# Linkage blocks involving multiple replicons organize *Borrelia burgdorferi* strain variation and influence virulence in Lyme disease

**DOI:** 10.1101/2025.01.29.635312

**Authors:** Rachel A. Laing, Michael J. Foster, M. Amine Hassani, Benjamin Kotzen, Alexander Arvanitis, Weihua Huang, Terrence Shea, Ali A. Djarou, Gordon C. Adams, Stephen F. Schaffner, Tjasa Cerar, Lisa Freimark, Eva Ruzic-Sabljic, Dionysios Liveris, Radha Iyer, Brenda Rosario Perez, Kathleen Fitch, Linden Hu, Robert Smith, Kurt D. Reed, John A. Branda, Allen C. Steere, Gary P. Wormser, Franc Strle, Pardis C. Sabeti, Ashlee M. Earl, Ira Schwartz, Klemen Strle, Jacob E. Lemieux

## Abstract

The many plasmids (commonly >20 per strain) of *Borrelia burgdorferi* (*Bb*) pose challenges for studies of Lyme disease. The genetic content of most plasmids cannot be resolved using short-read sequencing. We generated long-read assemblies (LRA) of 183 isolates of human-derived *Bb* and analyzed patterns of genome variation in the Lyme spirochete. LRAs confirm that populations consist of strongly-structured genotypes with nearly-clonal structure and tight-knit blocks of accessory genome elements. Notably, these patterns of linkage were statistical but not physical, with linkage blocks distributed across multiple plasmids. A consequence of this structure is that plasmid name and/or plasmid subtype does not capture strain-specific genetic content. We used network methods to characterize patterns of genetic linkage. We demonstrate that co-occurring gene networks, here termed genetic modules, are the fundamental unit of genome variation in *Bb* and designate genetic modules consisting of co-occurring genes. We linked genetic modules to *Bb* phenotype by identifying modules that influence dissemination in humans. This modular decomposition clarifies previously observed associations between strain and virulence. For example, virulent RST1/OspC type A strains are distinguished by the presence of virulence-associated modules containing gene content from lp28-1, lp56, and the chromosome along with the absence of gene content on lp28-1 and lp28-4 associated with localized disease. LRAs also demonstrate that the well-established statistical linkage between physical unlinked genetic markers (e.g. RST and OspC) is a general pattern among accessory genome elements. In summary, genetic modules containing genes linked across multiple replicons, rather than strain-defining plasmids, organize the *Bb* accessory genome and the strain-specific variation responsible for differences in human virulence.

**AUTHOR SUMMARY:** *Borrelia burgdorferi* (*Bb*) sensu lato causes Lyme disease, the most prevalent tick-borne disease in the Northern hemisphere. *Bb* strains are genetically heterogeneous, and strain variation is linked to differential severity and clinical manifestations. The bacterium possesses one linear chromosome plus numerous circular and linear plasmids; collectively these plasmids comprise ∼40% of its genome. Because these plasmids are difficult to assemble when sequenced using short-read methods, patterns of genome variation have been difficult to resolve, impeding studies of *Bb* population genetics and genotype-phenotype associations. In this study, we used long-read sequencing to assemble high quality genomes of human-derived strains of *Bb*. The resulting genomes reveal the structure of genetic variation across the complete genome. These analyses confirm the nearly clonal expansion among *Bb* genotypes, which are characterized by complex genomic rearrangements. An unexpected finding is that clade-specific genetic content is not well captured by plasmid identity and includes co-varying content across multiple plasmids. The fundamental unit of variation is therefore neither plasmids nor plasmid subtypes but instead linkage blocks of co-occurring genes. We identify these blocks using gene co-occurrence relationships and annotate them as genetic modules. Scoring each gene based on distribution over dissemination status, we find that the presence or absence of certain modules is associated with human dissemination. Overall, the findings provide a comprehensive analysis of *Bb* genetic variation across its genome, including in difficult-to-resolve plasmid sequences, and will facilitate the identification of the spirochete genes that influence virulence and distinct clinical manifestations.

## INTRODUCTION

Lyme disease is the most common tick-borne disease in North America, with an estimated 476,000 cases of Lyme disease diagnosed annually in the United States [1]. In humans, the clinical manifestations of Lyme disease are variable and progress in stages. Disease begins with inoculation of the causative spirochetes, *Borrelia burgdorferi* (*Bb*), into the skin by the bite of an infected tick, usually resulting in a skin lesion (erythema migrans). If untreated, *Bb* may spread to other sites resulting in disseminated disease and potentially leading to multiple erythema migrans skin lesions, meningitis, carditis, arthritis, and/or other complications [2–4]. Most patients respond well to appropriate antibiotic therapy, but 5-20% of patients experience persistent pain, neurocognitive, or fatigue symptoms after treatment, termed post-treatment Lyme disease syndrome (PTLDS), a condition for which there is no effective therapy [5–7].

The complexity of the symptoms and clinical signs of Lyme disease is shaped by host and microbial factors. Variants in human loci, including HLA, secretoglobin, TLR1, and LCE3, have been associated with the risk of Lyme disease or certain complications [8–11]. The importance of microbial genes is highlighted by the relationship between spirochete diversity and clinical manifestations. *Bb* comprise a diverse clade within the family *Borreliaceae*, individually labelled “genospecies” and collectively referred to as *Borrelia burgdorferi* sensu lato, Lyme borreliae, or the recently proposed genus nov *Borreliella* [12–14]. *B. garinii* and *B. bavariensis* are associated with Lyme neuroborreliosis; *B. burgdorferi* sensu stricto (*Bbss*, or here simply *Bb)* is associated with arthritis, and *B. afzelii* is associated with a chronic skin infection, acrodermatitis chronica atrophicans. Within *Bb,* strains may be further subdivided into genotypes based on a variety of methods including serotyping [15], 16S-23S intergenic ribosomal spacer typing (RST) [16], outer surface protein C sequencing (OspC) [17], multi-locus sequence typing (MLST) [18], and whole genome sequencing [19]. A consistent finding among typing studies is that certain genotypes of *Bb* are more likely than others to disseminate [20–23], more likely to cause Lyme arthritis [24], and are associated with greater inflammation [10,25–27]. Thus, *Bb* genetic diversity is linked to clinical manifestations in all branches of the spirochte tree and across phylogenetic scales; however, the individual genes underlying these associations have not been mapped in detail.

Genetic analysis of *Bb* strains has been challenging for two reasons. First, human isolates of *Bb* are difficult to obtain because they typically require an invasive diagnostic procedure (such as a skin biopsy) and specialized culture methods. Second, the *Bb* genome has a unique structure consisting of a single chromosome of approximately 910 kb and numerous plasmids (21 in the B31 reference strain, which has a total length of approximately 1500 kb) [28]. Approximately 40% of the gene content is encoded on multiple linear plasmids (named with the prefix lp) and circular plasmids (named with the prefix cp). Short-read sequencing cannot distinguish most plasmids due to their homology [29]. However, characterizing plasmids is critical because they are strain-variable [19,30,31] and encode determinants of infectivity [32]. Newer approaches from Oxford Nanopore Technologies (ONT) and Pacific Biosciences (PacBio) can produce longer sequence reads that resolve *Bb* plasmids [29,33].

In this study, we used long-read sequencing to characterize genome variation among human *Bb* isolates. We aimed for a substantial and representative dataset among human-invasive isolates in the United States. We characterized plasmid composition, patterns of genetic variation across the entire genome including plasmids, and analyzed associations of genetic content with dissemination in humans.

## RESULTS

### Long-read assemblies resolve the *Bb* chromosome and plasmid sequences

We sequenced and assembled 183 *Bb* strains (182 human isolates and 1 tick isolate), and sourced 17 non-overlapping sequences (6 human isolates and 11 tick isolates) from GenBank (Table S1-2) [34]. A subset of 146 genomes was previously analyzed by short-read sequencing [19]. Compared to short-read assemblies (SRAs), long-read assemblies (LRAs) yielded more complete replicons (Figure S1A), resulted in more contiguous assemblies as seen by higher N50s and fewer overall contigs (Figure S1B-C), despite no significant difference in number of unique genes predicted or total assembly length between sequencing approaches (Figure S1D-E).

We annotated LRAs for gene content and plasmid identity (Figure 1A). Plasmids were annotated using Pfam32 plasmid compatibility sequences (PF-32 genes) when present [29], or by BLAST nucleotide homology against reference plasmid sequences for replicons lacking a Pfam32 gene (cp9, lp5, and incomplete or fragmented contigs) (Methods). We estimated the frequency of plasmids across our sample overall (Figure 1B), as well as among major genotypes (Figure S2). We classified plasmids as either core (present in ≥95% of isolates) or accessory (<95% of isolates). Based on this criterion, the core plasmids are a 26 kilobase (kb) circular plasmid (cp26), a 54 kb linear plasmid (lp54), and a 17 kb linear plasmid (lp17). Other high-occurrence plasmids were lp36, lp28-3, lp28-4, and lp38, present in ≥75% of isolates (Figure 1A-B). In contrast, many plasmids were present only in certain genotypes. For example, cp32-4 and lp28-2 were more common among RST1 and OspC type A isolates, whereas cp32-2, lp28-5 and lp28-6 were more common among RST2 and OspC type K isolates (Figure S2).

**Figure 1:**
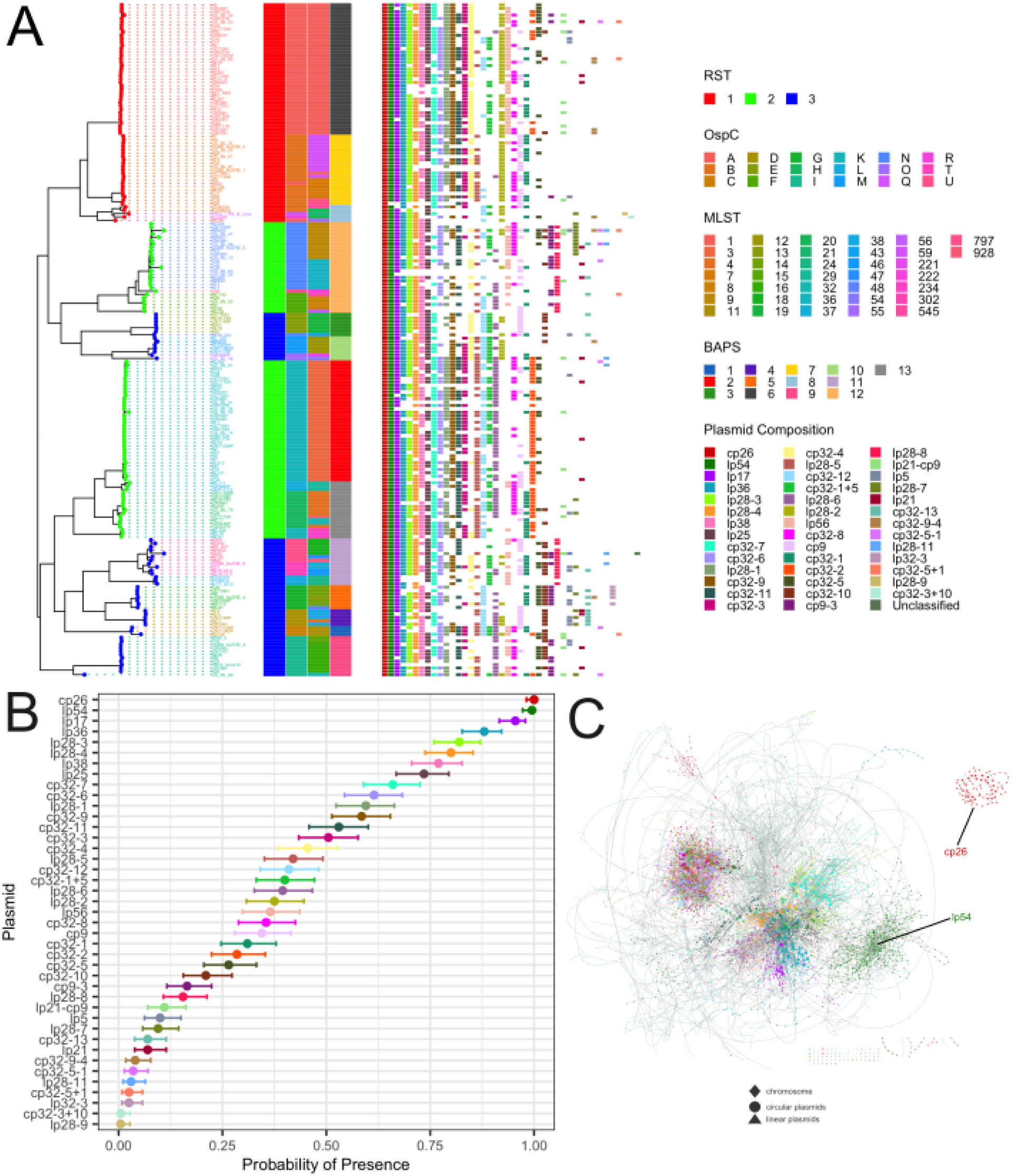
The phylogenetic structure of genetic markers and plasmid presence. (A) A phylogenetic tree assembled from core genome sequences is shown. Tips are colored by RST and tip labels are colored by OspC type. Triangles indicate samples on which RST was imputed. To the right of the tree, the results of genetic marker and cluster annotation from several typing schemes (RST, OspC, MLST, BAPS) are displayed. Plasmids are annotated by color and ordered by prevalence across the sample (n=200). (B) Clopper-Pearson 95% confidence intervals for plasmid presence according to our sample. (C) Pangenome graph generated by Panaroo as visualized using Cytoscape. Nodes are colored by their corresponding plasmid identity and positioned via Cytoscape’s Prefuse Force Directed Layout using the following parameters: (Iterations: 100, Default Spring Coefficient: 1E-4, Default Spring Length: 50, Default Node Mass: 3). Nodes are colored by encoding plasmid according to the same palette used in (A) and (B).

### Phylogenetic analysis reveals the structure of genetic variation in the *Bb* pangenome

We next analyzed patterns of gene content, focusing on overall genetic structure and the relationship between genes and plasmids. We constructed a pangenome by identifying ortholog groups and dividing them into core genes (ortholog groups shared by ≥95% isolates) and accessory genes (ortholog groups present in the sample but not in the core) using Panaroo [35]. The *Bb* pangenome consists of approximately 898 core ortholog groups and 5,120 accessory groups when paralogs are grouped together (Table S3). Separating paralogs into different groups, the core genome decreases to 862 gene groups, whereas the accessory genome expands to approximately 6,565 groups (Table S3).

A graph-based approach to pangenome construction, as provided by Panaroo [35], enabled us to quantify the sharing of genetic content between replicons (chromosome and plasmid sequences). In the pangenome assembly graph, cp26 resolves as a distinct cluster, and lp54 shows minimal connectivity with the pangenome assembly graph, indicating the genetic stability and functional integrity of these two plasmids (Figure 1C). The chromosome and other linear plasmids displayed greater connectivity, suggesting inter-replicons gene movements. In contrast, cp32 plasmids were densely connected to themselves (Figure 1C). These observations indicate that in general, with the exception of cp26 and lp54, plasmid type is not a reliable indicator of encoded genetic content. To assess sharing of genetic content using an alternate approach, we aligned each assembly to the B31 reference strain and assessed pairwise amino acid homology between the aggregated LRAs and the B31 reference. Similar to the pangenome graph, shared blocks of homology corroborated our observation that cp26 and lp54 form distinct genetic clusters, the frequent sharing of genetic material among linear plasmids and the chromosome ends, and the high homology among cp32 plasmids (Figure S3A-B). Taken together, these analyses indicate the genetic stability and functional integrity of cp26 and lp54 and further point to inter-replicon movement of genes in the *Bb* genome.

We next investigated relationships between gene presence, plasmid presence, and the phylogeny of isolates. To begin, we plotted the presence of ortholog groups by isolate, and annotated the encoding plasmid by color (Figure 2). As previously observed [19], sets of orthologs clustered along different branches of the phylogenetic tree. Annotation of the encoding plasmid revealed that clade-specific gene content was often, but not always, organized on one or more clade-specific plasmids. However, in several notable cases the patterns of ortholog groups were maintained but the encoding plasmids differed. For example, gene content from lp28-6 was found on lp28-2 in RST1 isolates (Figure 2). Additionally, plasmids with the same name encoded different genetic content at the clade-level, such as on lp38 and lp28-1 (Figure S4). Thus, although plasmid presence is correlated with gene content, the relationship is complex, with clade-specific gene content.

**Figure 2:**
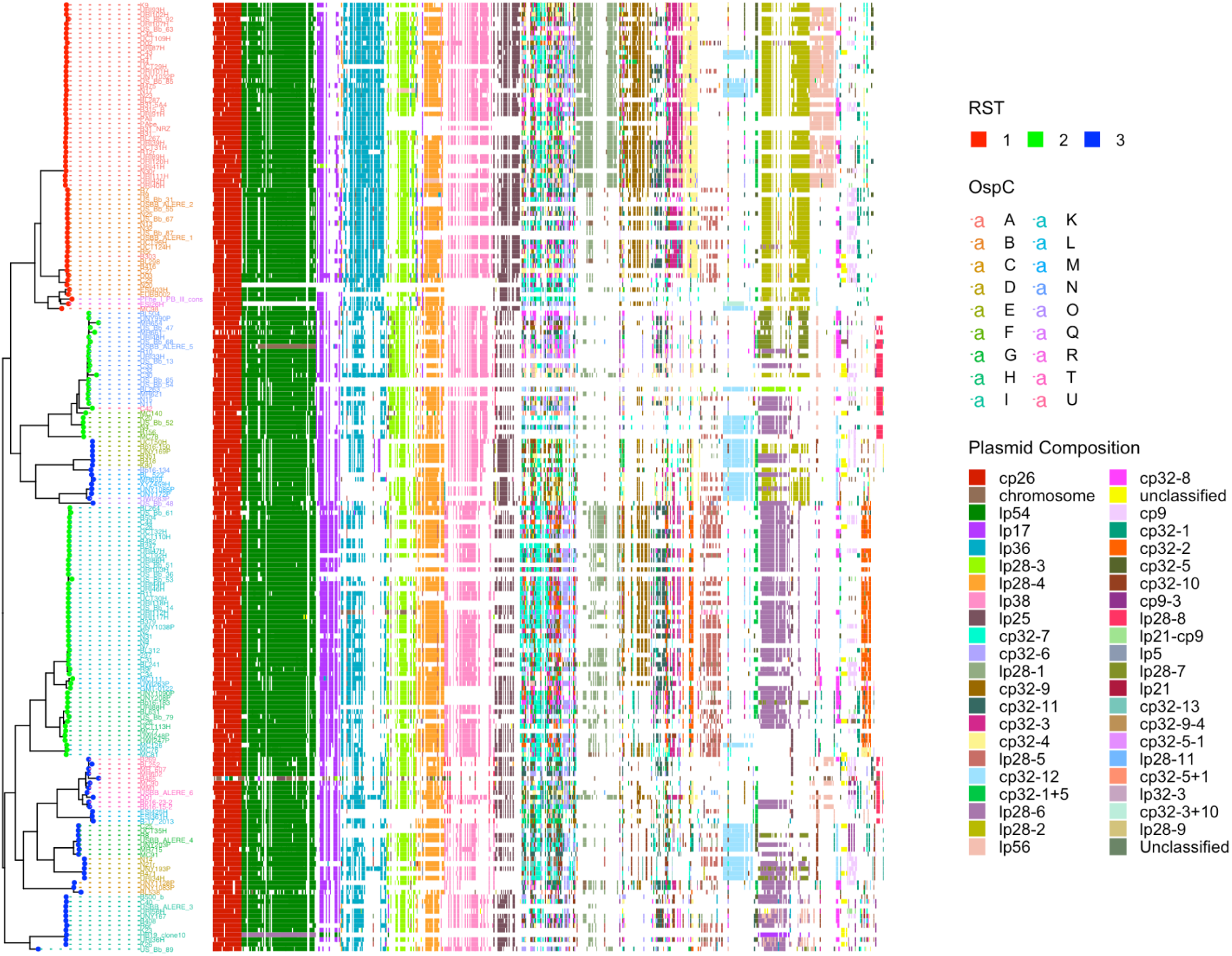
Phylogenetic structure and plasmid occupancy patterns of *Bb* orthologs. A phylogenetic tree assembled from core genome sequences is shown at left. Tips are colored by RST and tip labels are colored by OspC type. To the right of the phylogenetic tree, orthologs present in each isolate are shown in a matrix. Orthologs which were most commonly found on the chromosome were removed to emphasize patterns amongst replicons. For visual clarity, only orthologs present in at least 10% of isolates are shown. Columns of the matrix are grouped by best hit replicon, with replicons ordered by presence rates across our sample. The plasmid encoding the ortholog is annotated by color.

Given the importance of lipoproteins for *Bb* virulence [3,36,37], and their enrichment on plasmids [38], we analyzed patterns of lipoprotein variation in greater detail. Lipoproteins mirrored the broader pangenome pattern, with clade-specific gene content organized across multiple clade-specific plasmids (Figure S5A). Certain genotypes, particularly RST1/OspC type A strains, disseminate at higher rates and cause more severe disease [19–25,39]. These strains encoded a greater number of lipoproteins than other genotypes (Figure S7B), but plasmid count did not differ significantly between RST types (Figure S7C).

### Genomic similarity analysis reveals clonal expansion in *Bb*

To investigate the genetic structure of populations using near-complete genomes, we projected the isolates into two dimensions using principal component analysis of the pangenome via encoded ortholog groups (Figure 3B). This two-dimensional embedding revealed that RST1 isolates formed a mostly distinct subpopulation compared to RST2 and RST3 isolates (Figure 3B). RST2 and RST3 are polyphyletic, consistent with short-read phylogenies [19]. Apart from the clear separation of RST1 isolates, the population structure is complex, with different typing systems capturing different granularity (Figure 3B and Figure 1). To quantify similarity between *Bb* genomes, we computed pairwise average nucleotide identity (ANI) among the LRA-sequenced isolates and plotted these measurements alongside the core genome phylogeny (Figure 3A-B). The block structure of observed ortholog presence/absence was also seen using ANI. Co-occurrence blocks tended to correspond to OspC types. To quantify this, we assessed pairwise genetic diversity between all OspC types, where isolates of the same OspC types typically had a high degree of genomic similarity (Figure S8A). Comparing core and accessory genome divergence, there was an approximately linear relationship between accessory and core genome distance, but accessory genome variation was multi-modal at a given core genome divergence, indicative of episodic, marked changes in accessory genome content (Figure S8B).

**Figure 3:**
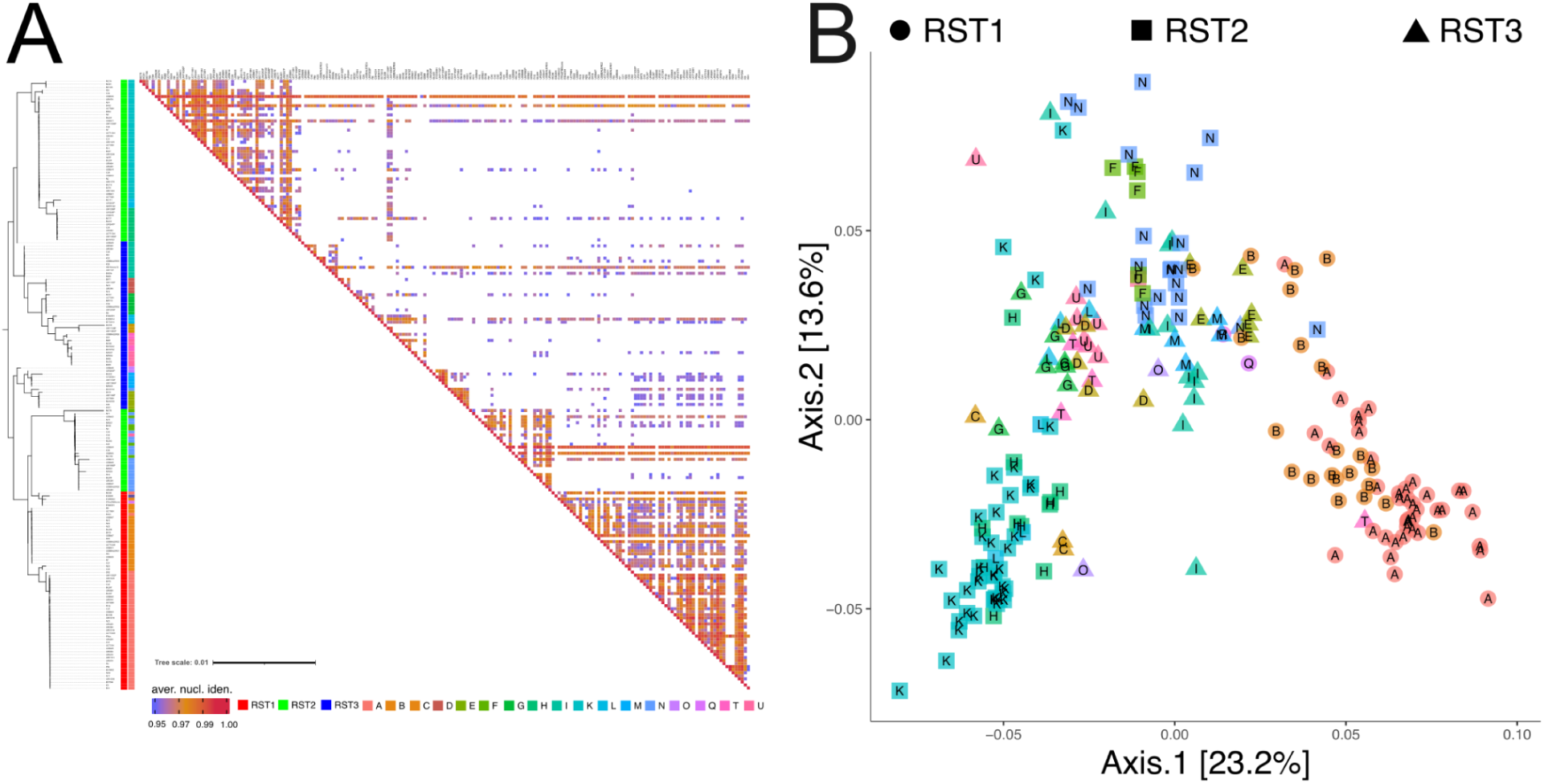
Genetic structure and patterns of divergence in the core and accessory *Bb* pangenome. (A) The phylogenetic tree constructed from core genome sequences is shown at left. Genetic markers and cluster membership are shown next to the phylogenetic tree. The heatmap shows the upper triangular portion of a matrix whose elements correspond to the pairwise calculation of full average nucleotide identity (ANI, cutoff >0.95). Each cell of the matrix is colored by ANI value. (B) Principal component analysis shows the projection of Sørensen distances for the collection of isolates. Marker shapes represent RST, and annotated letters indicate OspC type.

### Gene co-occurrence patterns define modules within the *Bb* genome

The patterns of accessory genome variation have a block structure indicating statistical linkage of accessory elements. This block pattern of gene presence (Figure 2) is maintained despite changes in the encoding plasmids, and patterns extend beyond a single replicon. To capture these patterns, we created correlation networks of accessory gene groups (i.e., not present in all isolates) and visualized their structure through clustering and module analysis.

We annotated networks using a colorspace mapping (Methods); for three genotypes (ribosomal spacer types), this corresponds to a triangle (Figure 4A). In the center (in white) lie the gene groups which are present in a vast majority, but not in all, of the sampled isolates. Universal across the genotypes, these are likely related to core functions. Conversely, at the corners lie the gene groups which are specific to singular RST types. These gene groups may be responsible for variations in clinical manifestations in patients. Gene groups shared between RST types are dispersed among the triangle, highlighting the evolutionary history and genetic exchanges within *Bb*. This representation reveals the full scope of connections within the *Bb* genome, from core genes in near-white to common RST-dominant accessory genes in red, green, and blue extending from that central region and finally to small clusters of low abundance peripheral genic content. Genes shared across multiple genotypes are present as transition regions between distinct subsections of the network. We created an analogous network among lipoproteins (Figure 4C), which mirrors the high complexity of the full network. In particular, the lipoprotein structure is characterized by several uncommon genes, highlighting the variation in potential determinants of virulence and infectivity. Overall, there is a core network of orthologs present in nearly all strains with subclusters that correspond to RST, indicating that *Bb* gene relationships exist in clusters, some of which are unique to genetic subtypes.

**Figure 4:**
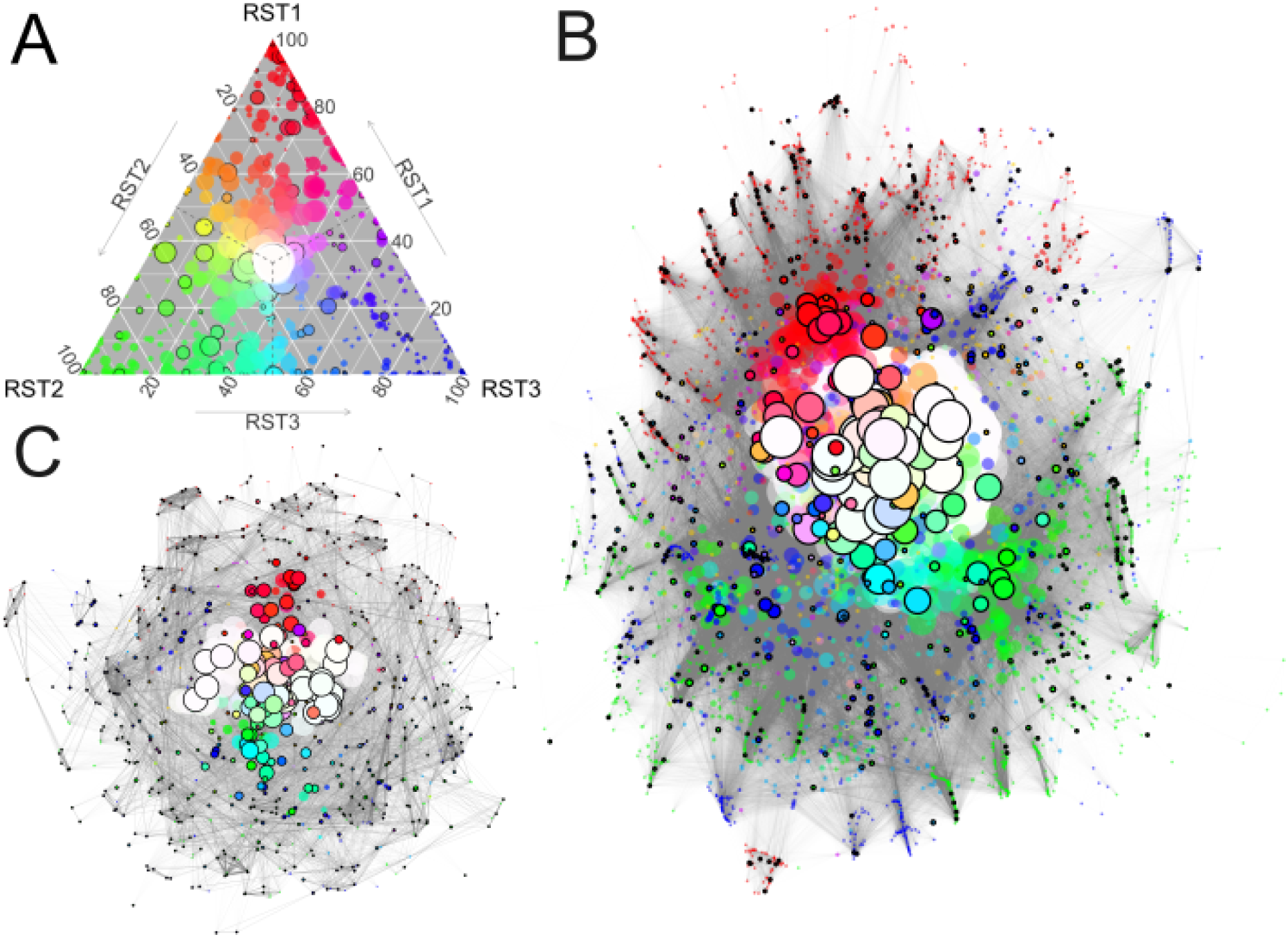
Analysis of the *Bb* pangenome correlation network reveals overarching patterns in underlying gene content. (A) Ternary plot of gene groups by RST types, with each point representing the normalized proportion of isolates within a specific RST type containing that gene group. Points are sized by gene frequency across the sample, and coloring is determined with a 3-dimensional RGB color cube mapping based on gene distribution among RST types. Surface lipoproteins are circled in black. For visibility, a small jitter has been applied to the points. This also doubles as an intuitive legend / graph for the network figures, where all of the same stylistic choices apply. (B) Positive correlation network for all gene groups. (C) Positive correlation network for gene groups associated with lipoproteins.

### Biological associations of genetic modules

The prominence of linkage blocks and their correlationships with phylogenetic clades highlighted the need for systematic tools to characterize these blocks and investigate their statistical properties and potential functions. We used a network approach to define modules of co-occurring genes. First, we created a subgraph of the full correlation network with genes present in two or more isolates and a minimum correlation threshold of 0.25, chosen to filter out trivial gene relationships whilst still preserving cross-genotype connections (Supplemental Methods). Then, maximum cliques were identified iteratively and plotted if they contained at least ten genes (Figure S9-10). The gene correlation heatmap confirms that genes within modules show strong patterns of statistical linkage with other genes in a given module. In some cases, negative correlations occur between modules, suggesting that modules may substitute for one another (Figure S11). Color coding the modules reveals that many were genotype specific. The top four modules by size are shown in Figure 5A. Phylogenetic trees on those gene subsets for Modules 1 and 2 are shown in Figure 5B. Module 1 (n=109) is RST1-dominant and contains genes from lp36, lp38, lp25, cp32-9, lp28-1, lp56, and more, representing a variety of linear and circular plasmids. Relative to the phylogeny, it is clear that these genes are primarily contained on the upper RST1 clade (composed of ospC type A), while the lower RST1 clade (composed of ospC type B) is missing the lp28-1 and lp56 sections. We note that the lp38 subset is also present on the upper RST2 clade and the upper and lower RST3 clades. Module 2 (n=86) is RST2 dominant and similarly features genes from lp38 and lp28-1, with contributions from lp28-5 and lp28-6 among others. These genes are attributed to the lower RST2 clades, with some degree of separation seen between the lower third of this subset to the rest. In this case, the lp28-6 genes are also seen partially in the upper RST2 clade, as well as the upper and lower RST3 clades. Module 3 (n=58) is shared across all RST types with genes that are exclusively found on lp54, whereas Module 14 (n=28) is RST3-dominant with a variety of encoding plasmids. In fact, all modules aside from Module 3, 5, and 38 span multiple plasmid families (where subtypes of lp28s and cp32s have been collapsed). In summary, modules are often genotype-specific, possess correlation patterns suggestive of functional relationships, and were frequently composed of genes encoded by different plasmids, indicating that statistical linkage is not mediated by physical linkage.

**Figure 5:**
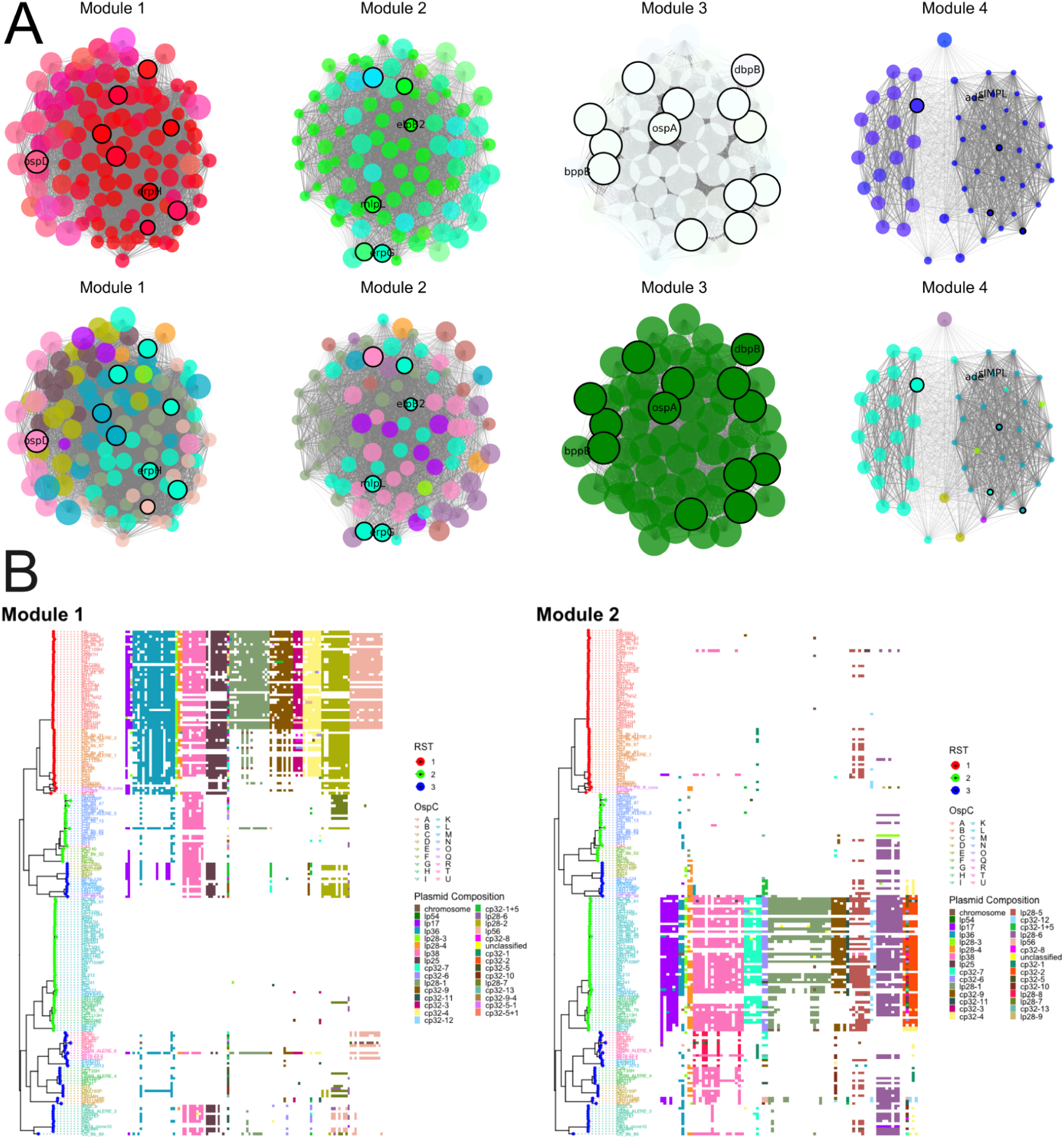
The *Bb* pangenome correlation network can be thresholded and partitioned into modules that uncover cross-plasmid gene group connections. (A) The largest four cliques of the thresholded graph with nodes colored by (i) distribution across RST and (ii) most commonly associated replicon. (B) Phylogenetic trees corresponding to Modules 1 and 2, providing additional context for these clusters.

The *Bb* accessory genome is enriched in surface-exposed lipoproteins, many of which are involved in interaction with the host. We annotated lipoproteins using computational identification of the lipoprotein leader sequence and investigated the relationship between lipoproteins and modules. Several modules were enriched in surface lipoproteins (e.g., Modules 13, 14, 26, 3, 19, 21, 7, and 27; p values < 0.05 by hypergeometric test (Table S4)). Certain modules also contain multiple variations of surface lipoprotein families (Table S4). From the set of enriched modules, Module 13, 14, and 27 have lipoproteins from the *erp* family, as well as *ospB*, *ospD*, and *mlp* respectively. Modules 3 and 21 both include *ospA* and *ospB*, with Module 3 also having *dbp* and Module 21 having *mlp*. Modules 26, 19, and 7 only include single lipoprotein families, namely *erp*, *mlp*, and *erp* respectively. Module 1, which is not statistically significant in lipoprotein presence, includes several lipoproteins that known to influence virulence in murine models with *mlp*, *erp*, and *vls*. Additionally, the lipoprotein composition of both Module 1 (an RST1-specific module) and Module 2 (an RST-2 specific module), include *mlp* and *erp* groups. A complete list of modules and their gene groups is provided in Table S5. Each module is assigned a primary RST type based on the composition of genes within it, resulting in 17 RST1 modules, 17 RST2 modules, 12 RST3 modules, and 3 core modules (where a majority of genes contained were present in at least 95% of isolates within each RST group). Overall, certain genotype-specific modules contained known determinants of virulence, and some were enriched in surface lipoproteins.

We next sought to identify modules associated with dissemination by linking clinical characteristics of patients from whom the isolates were recovered to the annotated modules. Isolates were labeled as disseminated if *Bb* was detected in blood, multiple erythema migrans lesions were observed, or the patient presented with CNS disease (Methods). Out of the 188 human isolates, 78 were annotated as disseminated, 76 as localized, and the remaining 34 did not have this information available. Contingency tables of RST and OspC types versus dissemination confirm previously observed associations[20,22,23] between these markers and dissemination status (Table S6). For each ortholog group, we calculated presence proportions within both the disseminated and localized classifications. We then took the difference between these values to assess the tendencies of each gene group, and incorporated that into the module visualization (Figure 6A). Gene groups within the same modules tended to have similar propensities towards dissemination. Difference scores were averaged within each module and plotted in numeric order (Figure 6B), with lipoprotein-enriched modules (as discussed in the prior section) highlighted. Module 22 is definitively the top candidate for strongest weighting towards disseminated isolates, and Module 28 represents the opposite, favoring localized isolates. When modules are plotted alongside phylogenetic trees (Figure 6C), the genotype-distribution is apparent: Both modules consist of gene groups that are primarily specific to RST1 strains. However, Module 22 contains gene groups from the upper RST1 clade, all of which are also OspC Type A, while Module 28 represents the lower RST1 clade which are primarily non-OspC Type A isolates. These modules consist of genes encoded on multiple replicons, particularly lp28-1, lp56, and the chromosome (Figure 6C). These patterns not only exemplify the linkage structure of the *Bb* accessory genome variation but also suggest that the unique virulence of RST1/OspC A strains is a consequence of variable genetic content rather than the strain-specific presence of certain plasmids.

**Figure 6:**
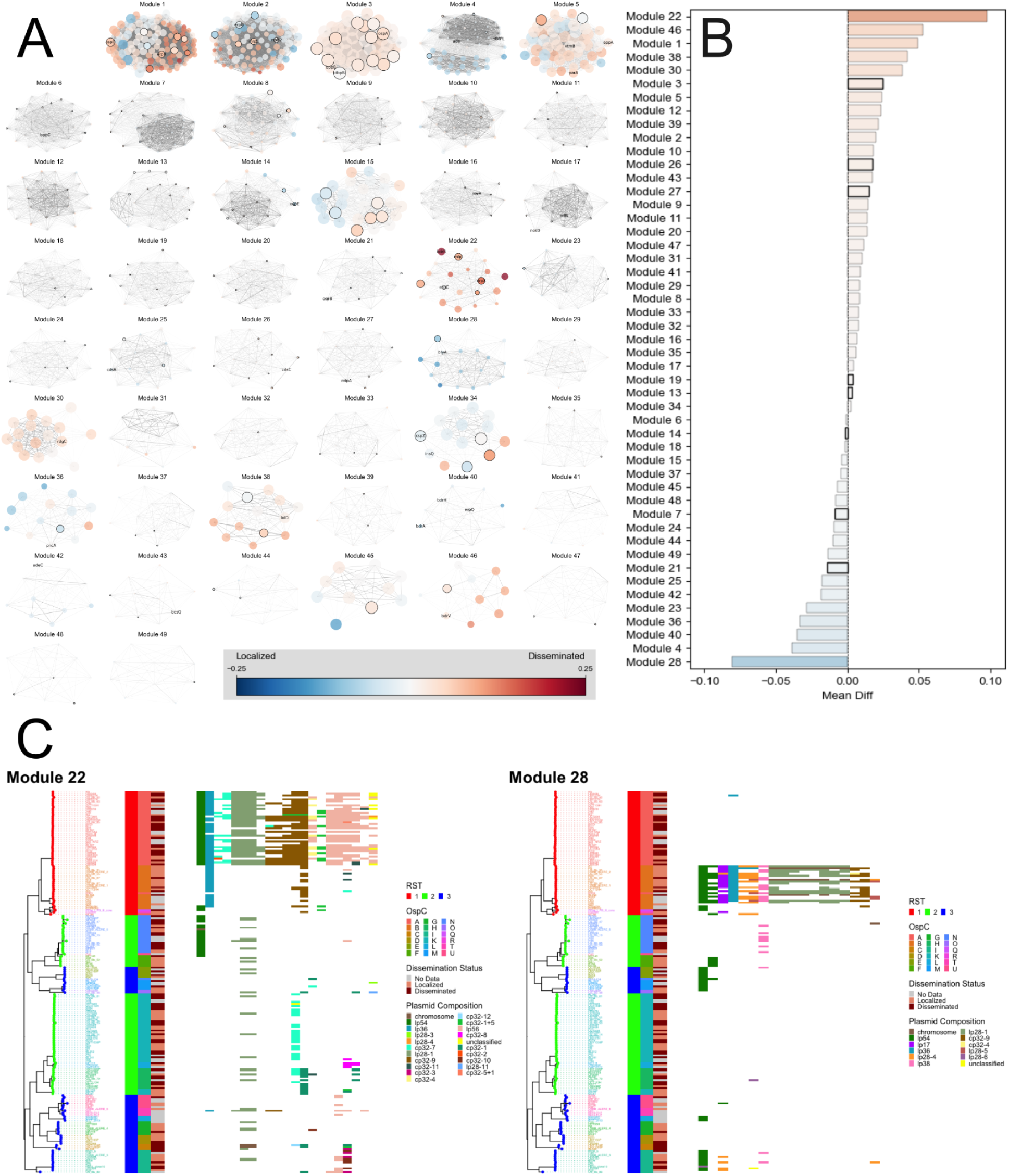
Modules of the *Bb* pangenome demonstrate various levels of association with dissemination. (A) Cliques were found iteratively upon a subgraph of the full correlation network where genes present in only one isolate were dropped and the minimum correlation threshold was set to 0.25. For clarity, only cliques with 10 or more nodes are displayed. Node colors are determined based on the difference between a gene’s presence level in disseminated versus localized isolates; isolates with unavailable clinical metadata were dropped from this analysis. Surface lipoproteins are circled in black, and certain gene groups with sufficiently short names are annotated. (B) Module-specific average weighting towards disseminated or localized isolates. (C) Phylogenetic trees corresponding to Modules 22 and 28.

## DISCUSSION

We present a comprehensive analysis of the population genomic variation of *Bb* isolates obtained primarily from North American/US patients with Lyme disease based on long-read sequencing, which effectively resolves plasmid-rich *Bb* genomes. Our results confirm existing observations about the *Bb* population structure, while offering new insight into the structure of *Bb* evolution, genetic diversity, and virulence.

LRAs provide a more complete picture of *Bb* genomes and population structure than has been previously available from short read analyses. The insights revealed LRAs both confirm several observations from single-locus and short-read studies and reveal new ones. We confirm that human-infectious *Bb* populations are strongly structured, which is particularly notable in Figures 2 and 3. This is consistent with the results of multiple decades of previous studies including those that have used serotyping [15,40], a variety of genetic markers (RST [16], OspC [17], MLST [18,39]), and short-read sequencing [19]. RST1 is monophyletic but RST2 and 3 are phylogenetically mixed. Among RSTs, RST3 appeared to be genetically heterogeneous, consistent with previous observations [24–26]. OspC types possess a high genetic similarity of most OspC types (Figure 3B), indicative of clonal expansion within these types.

Other investigators studying *Bb* have reported chromosomes of variable length, usually between 904 and 932 kb [28,41–47] with which our findings are consistent. Our chromosomal contigs show evidence of fusion with various linear plasmids as described elsewhere [30]. We also observed genetic material from various other linear and circular plasmids being appended to the chromosome (Figure S3). This has previously been observed with regard to the variable presence of BB0844 at the chromosomal right end [48], and proposed mechanisms mediated by the telomere resolvase ResT [49]. However, the genetic isolation of cp26 and lp54 is notably revealed by whole genome comparisons of pangenome contiguity and homology.

Long reads also provide new insight in several areas. First, this analysis provides the largest survey today of the patterns of plasmid presence among *Bb* isolates. Cp26 and lp54 are ubiquitous among all sequenced *Bb* to date [50,51], and are essential for the infection cycle [52,53]. The frequency of all plasmids is provided in overall Figure 1B and by genotype in Figure S2 and will be a resource to other investigators.

Second, LRAs also reveal the structure of populations in the population in the accessory genome. Previous studies using single genetic markers [54] and whole genome sequencing with short-reads [19] identified a high linkage disequilibrium between markers of different replicons in *Bb* genomes. LRAs reveal this statistical linkage without physical linkage to be a general pattern of *Bb* genomes. These findings are consistent with a near-clonal structure among *Bb* genotypes. Accessory divergence at a given core genome divergence, along complex yet phylogenetically consistent relationships, imply that multiple, complex evolutionary changes have occurred and are maintained in *Bb* populations. These observations collectively suggest that complex rearrangements of the accessory genome have occurred and then undergone clonal expansion.

Third, linkage among genetic elements involves multiple, physically-unlike plasmids (Figure 2). This implies that evolutionary forces beyond physical linkage maintain this structure. Supporting this, patterns of co-occurrence persist even when the encoding plasmid changes. These observations suggest that functional relationships have developed among linked orthologs.

A major goal of this analysis was to resolve plasmids using long-read sequencing. This approach was technically successful. Yet, somewhat surprisingly, knowing which plasmids were present in a given isolate provided little insight into population structure or virulence. For example, knowing that lp38 or lp28-1 is present does not predict which genes are encoded in a given strain (Figure S4). An updated plasmid nomenclature that designates lineage-specific plasmids with a unique identifier could increase the information captured by plasmid name. Invasive genotypes also do not appear to possess larger numbers of plasmids (Figure S7A-C).

Improved plasmid nomenclature, however, would not make plasmids fully informative for isolate genotype or phenotype because linked genetic content is encoded on multiple different plasmids (e.g. Figure 6C). While genotypes with higher virulence did not differ in their plasmid content, invasive genotypes did have greater numbers of lipoproteins (Figure S7B). This observation, consistent with previous analyses using short read data[19], also supports the importance of gene content rather than plasmid content for determining virulence. Thus, more so than plasmids, patterns of gene co-occurrence appear to be the fundamental unit of genome variation in the accessory genome of *Bb.* Plasmids may be better understood as flexible vessels for underlying genetic content rather than determinants of bacterial properties such as virulence and infectivity themselves.

We used clique-based analysis to identify the genes that likely comprise these functional networks (Figure 4). By capturing the positive- and negative-linkage patterns between genes, modules provide insight into the genetic differences between strains and may help guide laboratory investigations into determinants of virulence and other phenotypes. The robust linkage structure and the negative correlations between some modules suggest that some modules can substitute for one another and supports functional relationships for some of these network modules. For example RST1 strains, particularly the subset marked by OspC type A, are more virulent than other strains [20–23,25]. While most RST1 strains possess module 1 (with gene content encoded on lp36, lp38, lp28-1, and lp56), associated with dissemination, only OspC type A strains possess module 22 (most strongly associated with dissemination, with gene content encoded on lp28-1, lp56, and the chromosome) and lack module 28 (associated with localized disease, with gene content on lp28-4 and lp28-1). These modules include allelic variants of known virulence determinants such as OspC [55,56] and DbpA [57], as well as numerous unannotated genes that are candidates for future study.

Previous genetic association studies have implicated lineage-level effects but failed to capture locus-level effects because of strong linkage blocks [19]. Modules effectively offer a granular way to identify and characterize lineage-level effects. The causal structure of these associations needs to be explored further in future work. It remains unclear whether virulence and other are controlled by a small number of causal loci with many linked non-contributory loci, or if causal effects are polygenic, with many genes within a module working together to promote virulence.

This study has several limitations. The sampling was non-random and weighted toward North American clinical isolates. Rare OspC types are represented by one or only a few sequences, so any pertinent conclusions should be interpreted within this context. However, the distribution of genotypes in our study is representative of isolates obtained from patients, which increases the clinical relevance of these analyses. We also note that amongst our OspC type B isolates, 4 were disseminated, 12 were localized, and 6 had no data, which is an undercount compared to a prior cohort where slightly over 50% of OspC type B isolates were disseminated [22]. Sampling differences may be relevant to our findings regarding modules associated with dissemination. Furthermore, infections caused by a virulent strain that are detected and treated early would be considered localized despite having the potential to cause disseminated illness. Isolates were cultured and, despite attempts to minimize spontaneous plasmid loss by limiting passaging, may have lost plasmids during culture adaptation or library preparation [58,59]. In addition, although we generated assemblies by multiple different methods, we were not able to complete the genomes fully for certain highly homologous plasmids. Finally, given the complexity of *Bb* genomes, assembly errors are possible. Long-read assemblers struggle with short plasmids, often requiring special considerations to be taken into account [60]. The phylogenetic consistency of patterns across multiple sources of isolates, as shown in Figure 2, including those sequenced for this study and those from published assemblies, suggests that misassemblies and plasmid loss are not a major source of errors. However, due to the possibility of misassemblies, we consider these assemblies near-complete and not base-perfect.

In conclusion, this study provides the first detailed analysis of plasmid sequence diversity and genetic structure among *Bb*. The genomes and analyses presented here complement a recent report [61] on complete *Bbsl* genomes, as systematic surveys of *Bb* evolution using LRAs confirm and expand on observations from previous short-read analyses [19,34,62–64] and earlier analyses of plasmid diversity [31,50,65]. In particular, LRAs reveal the ways in which plasmids provide a mechanism for strain-specific inheritance of accessory genome elements and reveal the complex interchanges that occur among plasmids, while genetic networks remain relatively stable. This high-resolution analysis of the complete *Bb* pangenome provides new insights into the fundamental organizing principles of *Bb* genomes, defines a linkage between modular gene content and human virulence, and provides a framework for identifying *Bb* genes associated with distinct clinical manifestations, organ tropism, and other phenotypes.

## METHODS

### PacBio assemblies

Genomic DNA was extracted using the QIAamp DNA kit (Qiagen). The sequencing library was prepared using the SMRTbell template preparation kit (Pacific Biosciences). Sequencing was performed in the RSII SMRT sequencing system. After sequencing, reads were trimmed, filtered, and assembled using the Hierarchical Genome Assembly Pipeline (HGAP). Resulting contigs were trimmed for linear plasmids and the chromosome, and circularized for circular plasmids. All the contigs were manually polished and corrected by BWA alignment and mapping [66] and Integrative Genomics Viewer (IGV) [67] visualization using the previously generated Illumina short-reads [19].

### Oxford Nanopore Technologies (ONT) assemblies

We performed two different assembly methods: hybrid (i.e. long-read scaffolds polished with short-reads) and ONT long-read assemblies. Hybrid assemblies were generated as previously described [68]. Briefly, samples were barcoded using the Oxford Nanopore Rapid Barcoding kit SQK-RBK004 and run in batches of six samples per flow cell on a GridIon sequencer (Oxford Nanopore Technologies Ltd, Science Park, UK). ONT reads were demultiplexed using Deepbinner v0.2.0 [69], adapters were trimmed using Porechop v0.2.3 [70], and subsampled to ∼50× depth. Illumina reads were trimmed of adapters using Trim Galore (v0.5.0) and subsampled to ∼100× depth. Two Unicycler v0.4.4 [71] (with default settings) hybrid assemblies were generated for each sample, one assembly combining the Illumina 100× dataset with the 50× subsampled ONT dataset and another assembly combining the Illumina 100× dataset with the full set of ONT reads (if >50×). ONT reads were aligned to Unicycler contigs using minimap2 v2.15 [72]. Illumina reads were aligned to Unicycler contigs using BWA MEM (v0.7.17) [66], and the resulting alignments were input to Pilon v1.23 [73] for assembly polishing. For non-hybrid assemblies, demultiplexed raw BAMs were converted to fastqs using samtools fastq v1.22.2 [74] and then run through the Theiaprok Public Health Bioinformatics (PHB) pipeline v3.0.1 [75].

### For long-read ONT assembly

an additional 121 isolates were sequenced using the 24 and 96 barcode versions of the Oxford Nanopore native barcoding kit SQK-NBD114-24 and SQK-NBD114-96, respectively. These genomes were sequenced on a PromethION P2i for 72 hours and basecalled with Dorado using the SUP model (Oxford Nanopore Technologies Ltd, Science Park, UK). Raw reads were trimmed and filtered with Chopper [76], cleaned by taxonomic filtering with Kraken2 v2.1.6 [77], and assembled with Autocycler v0.5.2 [78]. Assembly polishing was performed with Dorado and final assembly graphs were cleaned using internally developed tools and visualized in BandageNG [79].

### Assembly quality control

All assembled contigs were validated to be free of contamination using Kraken2 using the “standard” database configuration [77]. Resulting assemblies were then analyzed using QUAST v5.2 [80] to generate assembly metrics with BioMark [81] for gene prediction counts. *Bb* B31 (GenBank assembly ASM868v2, NCBI RefSeq assembly GCF_000008685.2) was used as reference for preliminary determination of misassembly and relative genome content. To select the best assembly for each isolate, QUAST v5.2.0 [80], BUSCO v5.8.0 [82], and CheckM2 v1.1.0 [83] were run on the outputs of the Theiaprok and Autocycler pipelines. Assemblies with superior completeness, N50, and contig count, were retained.

### Genotyping and population structure analyses

For the determination of OspC typing, a BLAST database was constructed using sequences obtained from NCBI [84]. Each assembly was annotated via Bakta (v1.10.0) [85] and OspC sequences were aligned against this database. Alignments with highest identity were used as the OspC allele type and assemblies with multiple OspC alleles were determined to be contaminated and were dropped from further analysis. For determination of MLST type, we used mlst_check with the default databases bundled in the docker container provided by Sanger Pathogens [86]. Each assembly was then typed and the resulting output table merged with existing metadata. RST, MLST, and typing information was obtained from a published study [19] and studies reported therein. The hierarchical Bayesian Analysis of Population Structure (BAPS) was implemented in R using 2 levels of clustering and 9 initial clusters. The probability of cluster assignment was calculated for each genome [87]. Genomic similarity between the genomes was computed using PyANI program [88] implemented in Anvio (v8) and displayed full average nucleotide identity scores with a cutoff of 0.95 [89].

### Assembly annotation and pangenome construction

Generated assemblies were annotated using Bakta v1.11.0 [85]. The resulting gff3 annotations were then fed into the Core Gene SNP PHB pipeline v4.1.0, which we outfitted with Panaroo v1.7.0 for pangenome construction [78,39]. We ran panaroo with the following parameters: (--clean-mode strict -a core --aligner mafft --core_threshold 0.95 --remove-invalid-genes) both with and without (--merge_paralogs) (to examine the influence of paralog handling on the final pangenome. The pangenomic references generated were annotated with Bakta. We also predicted lipoproteins from the pangenome using an internally developed Python implementation of the SpLip algorithm [90] available at https://github.com/mjfos2r/BbLipoPredict that classifies putative signal peptides. Sørensen and Jaccard distances were computed using gene cluster presence/absence provided by Anvio pangenome analysis [89] implemented in the “vegan” R package [91]. Pairwise cophenetic distances were computed from the phylogenetic tree of core single gene clusters using the “stats” R package.

### Tree construction

The core genome alignment of panaroo was used to generate phylogenetic trees using IQTree v1.6.7 with a general time reversible model on nucleotide alignments [92]. Additionally, Geneious Prime 2025.0.3 (https://www.geneious.com) was used to generate various trees to assist manual review.

### Plasmid annotation

When possible, contigs were named using their PF-32 plasmid compatibility gene by aligning the contig to a database of PF-32 genes from GenBank and previous work [19]. If no PF-32 sequence was present, contigs were named according to BLAST homology with known plasmids from GenBank (BLAST v2.16.0). Contigs shorter than 1kb were considered too fragmented to reasonably annotate and were left unannotated. For disagreement between methods, manual review of BLAST homology and available annotations were used to resolve classifications to their most probable call.

### Lipoprotein identification

We identified lipoproteins by taking the union of those from Dowdell et al. [38] and our internal lipoprotein annotation program BbLipoPredict (available at https://github.com/mjfos2r/BbLipoPredict). Surface lipoproteins were further designated using the annotations previously published (S, P-IM, P-OM) [38], and additionally all gene groups relating to *osp*, *mlp*, *erp*, *dbp*, and *vls* were classified as surface lipoproteins [93].

### Phylogenetic visualization

Tree figures were created using R version 4.3.0 (https://www.R-project.org/), with the packages ggplot2, ggtree, ggtreeExtra, and tidytree [94–97]. All trees are midpoint rooted and formatted with the standard rectangular layout. Metadata columns are affixed to the tree object, then plotted as separate layers. We also provide interactive versions of the phylogenetic trees from Figure 2 and Figure 6 using the shiny package [98], which can be downloaded from our GitHub repository and run locally.

### Sankey homology calculation

All genomes were aligned with MMSeqs2 [99] wrapped by pygenomeviz (v1.4.2, pgv-mmseqs) (https://github.com/moshi4/pyGenomeViz), which performed alignment of translated CDS between each assembly and B31 to quantify the percent identity of amino acid sequences.

### Graph theory

We explored the relationships between genes using graph theory concepts and NetworkX v2.7.1 [100] in Python 3.11.3. Gene analysis was performed at the gene group level using presence/absence data from pangenome analysis. For all pairwise combinations of non-invariant gene groups, Pearson’s correlation was calculated. These correlations were then displayed via network format by representing the sign and strength of the correlation through the color and thickness of the edge between the two nodes. Further details on graph theory definitions and node color-mapping can be found in the supplement.

We utilized the correlation network to identify modular structures in the genome. First, all genes present in only 1 isolate were dropped and the network was thresholded to correlations in excess of 0.25. Then, an iterative algorithm was implemented to find the maximum clique, drop the involved nodes, and repeat on the resulting subgraph. This yields independent, non-overlapping, positively-correlated modules of gene groups, which may have functional links.

## DATA SUMMARY

All code used in this work is available at https://github.com/broadinstitute/Bb_pangenome. All genomes used in this work are detailed in Table S1 and are publicly available through NCBI/ENA. In addition to deposition within NCBI/ENA, all genomes and their annotations are available within the aforementioned repository. Clinical metadata on archived isolates were accessed for research purposes 9/1/2026. Coded specimens were used and patients were not identifiable to the authors for the analyses conducted.

## ACKNOWLEDGMENTS

We gratefully acknowledge the investigators who generated and submitted *Bb* genome data to GenBank.

