## SupplementaryMaterial for "Linkage blocks involving multiple replicons organize *Borrelia burgdorferi* strain variation and influence virulence in Lyme disease"

### Supplemental Methods

#### Key Definitions

A **graph** G = (V, E) consists of a set of vertices V and a set of edges E.

A graph $G_{sub} = (V_{sub}, E_{sub})$ is a **subgraph** of G if its vertices and edges are subsets of another graph, i.e. $V_{sub} \subseteq V$ and $E_{sub} \subseteq E$.

A graph $G_{ind} = (V_{ind}, E_{ind})$ is an **induced subgraph** of G if $V_{ind} \subseteq V$ and $E_{ind}$ contains every possible edge from E between the vertices in $V_{ind}$.

Two vertices are **adjacent** if they are joined by an edge.

A **clique** is a subset of vertices from a graph G such that every pair of distinct vertices in the clique is adjacent.

A **maximum clique** is the largest possible clique in a graph G.

**Fruchterman-Reingold Layout:** The Fruchterman-Reingold layout is a force-directed algorithm for the visualization of undirected graphs. Under this regime, highly linked nodes cluster together via attractive forces whilst other nodes are repelled [101]. In the genomic setting, clustering from a weighted correlation network can indicate plasmids or genes with similar expression patterns, as well as illuminate the behavior of accessory components [102].

**3-Dimensional Color Mapping:** We introduce a 3-dimensional color mapping scheme for network analysis of plasmids and genes in *Borrelia*. Taking advantage of the numerical correspondence between the three RST types and the three RGB values in uniquely defining colors, we assign RST1 to be red (RGB=(255,0,0)), RST2 to be green (RGB=(0,255,0)), and RST3 to be blue (RGB=(0,0,255)). Then, for each node (gene), the color is determined by the normalized proportion of isolates containing that node within each RST type. This can be represented by the following set of equations.

$$\mathrm{RST}[i]_{[\mathrm{gene}]} = \frac{\mathrm{RST}[i] \text{ isolates with [gene]}}{\text{Total number of } \mathrm{RST}[i] \text{ isolates}} \text{ for } i = 1, 2, 3$$

$$m_{[\mathrm{gene}]} = \max(\mathrm{RST1}_{[\mathrm{gene}]}, \mathrm{RST2}_{[\mathrm{gene}]}, \mathrm{RST3}_{[\mathrm{gene}]})$$

$$(R, G, B)_{[\mathrm{gene}]} = \left( \frac{\mathrm{RST1}_{[\mathrm{gene}]}}{m_{[\mathrm{gene}]}}, \frac{\mathrm{RST2}_{[\mathrm{gene}]}}{m_{[\mathrm{gene}]}}, \frac{\mathrm{RST3}_{[\mathrm{gene}]}}{m_{[\mathrm{gene}]}} \right)$$

Consider the simple example of Gene A present in 4/10 RST1 isolates, 1/5 RST2 isolates, and 2/8 RST3 isolates. This would lead to an RGB tuple of (0.4, 0.2, 0.25). Then, normalizing based on the maximum tuple value would yield (1, 0.5, 0.625), allowing movement along the most saturated three faces of the color cube. A representation of this process is depicted below, with arrows representing movement in the direction of decreased prevalence along a genotype. This mapping allows us to easily visualize which nodes are dominated by a certain RST type, as well as determine the RST combinations of any nodes shared between multiple types.

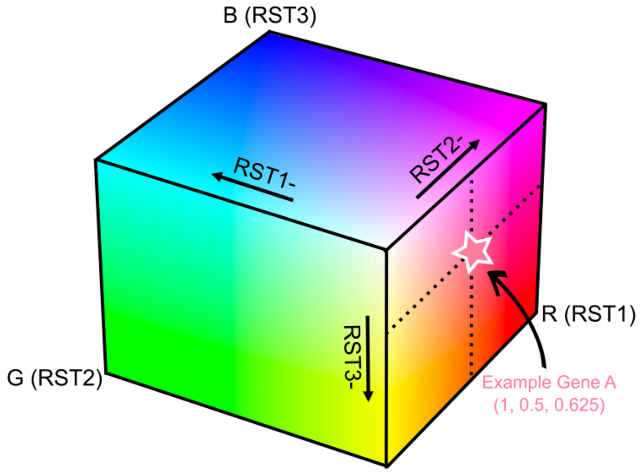


**Clique Analysis:** We partition the network into modules by iteratively identifying the largest clique, removing the involved nodes, and repeating this process on the resulting subgraph. This process is done on a thresholded positive correlation network, where all edges weaker than a correlation of 0.25 have been dropped. Correlation thresholds of 0 and 0.5 were also tested, with the former including too many insignificant gene relationships and the latter resulting in primarily small cliques and cutting out valuable cross genotype patterns that are crucial towards understanding the genetic structure of the *Bb* genome.

### Supplemental Figures

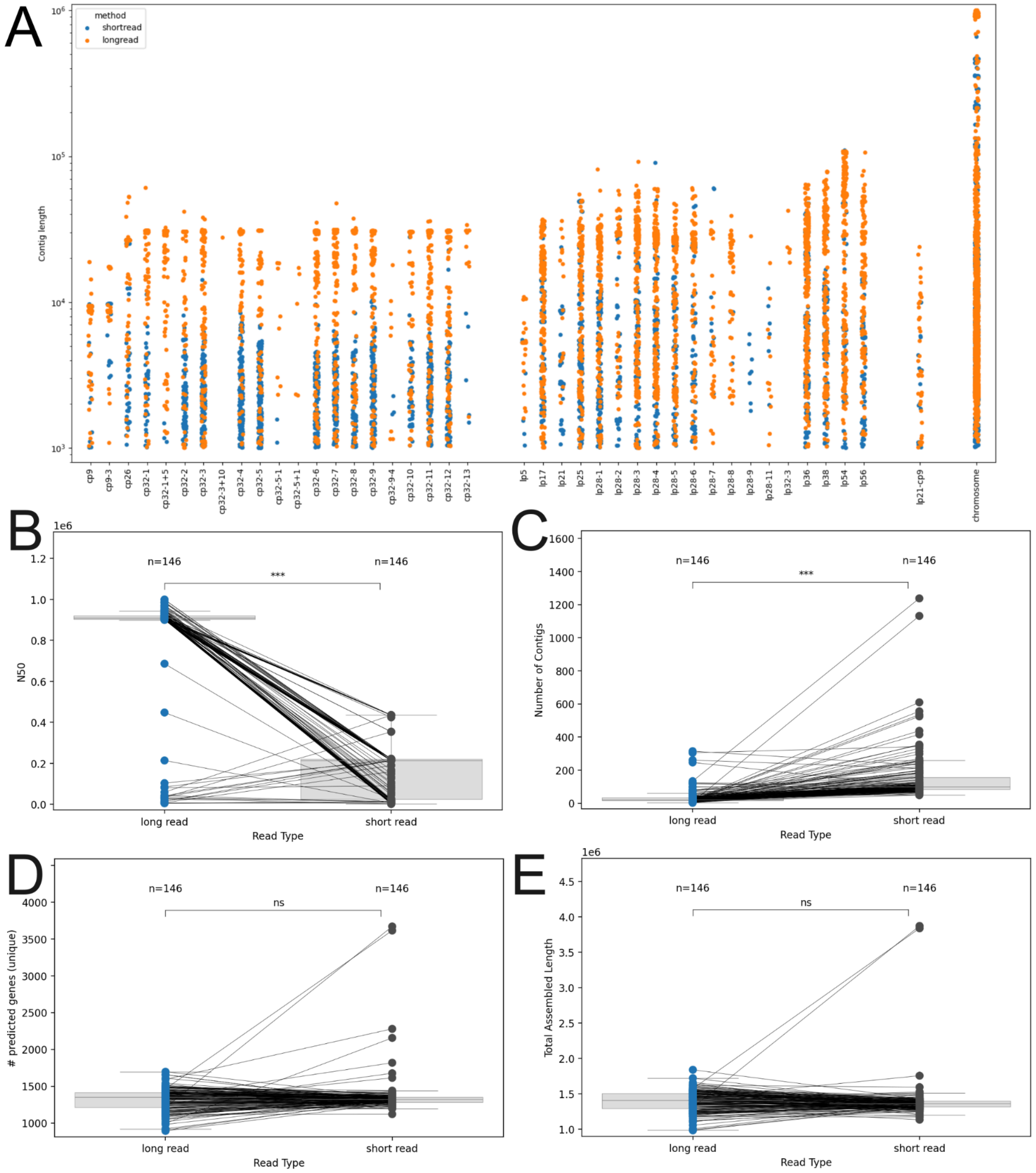


**Figure S1: Assembly quality comparisons between LRAs and SRAs.** (A) Jitterplot on the left shows the distribution of contig lengths over replicons, with LRAs marked in orange and SRAs marked in blue. Replicons are grouped by structure, with circular plasmids, linear plasmids, fusion plasmids, and the chromosome plotted from left to right. 146 assemblies that were also analyzed in a previous report [17] were considered in this analysis. (B) N50 of LRAs and SRAs with statistical significance shown (paired t-test; *** indicates p<0.01). (C) Number of contigs of LRAs and SRAs with statistical significance shown (paired t-test; *** indicates p<0.01). (D) Number of unique predicted genes in LRAs and SRAs with statistical significance shown (paired t-test; not significant) (E) Total assembled length of LRAs and SRAs with statistical significance shown (paired t-test; not significant).

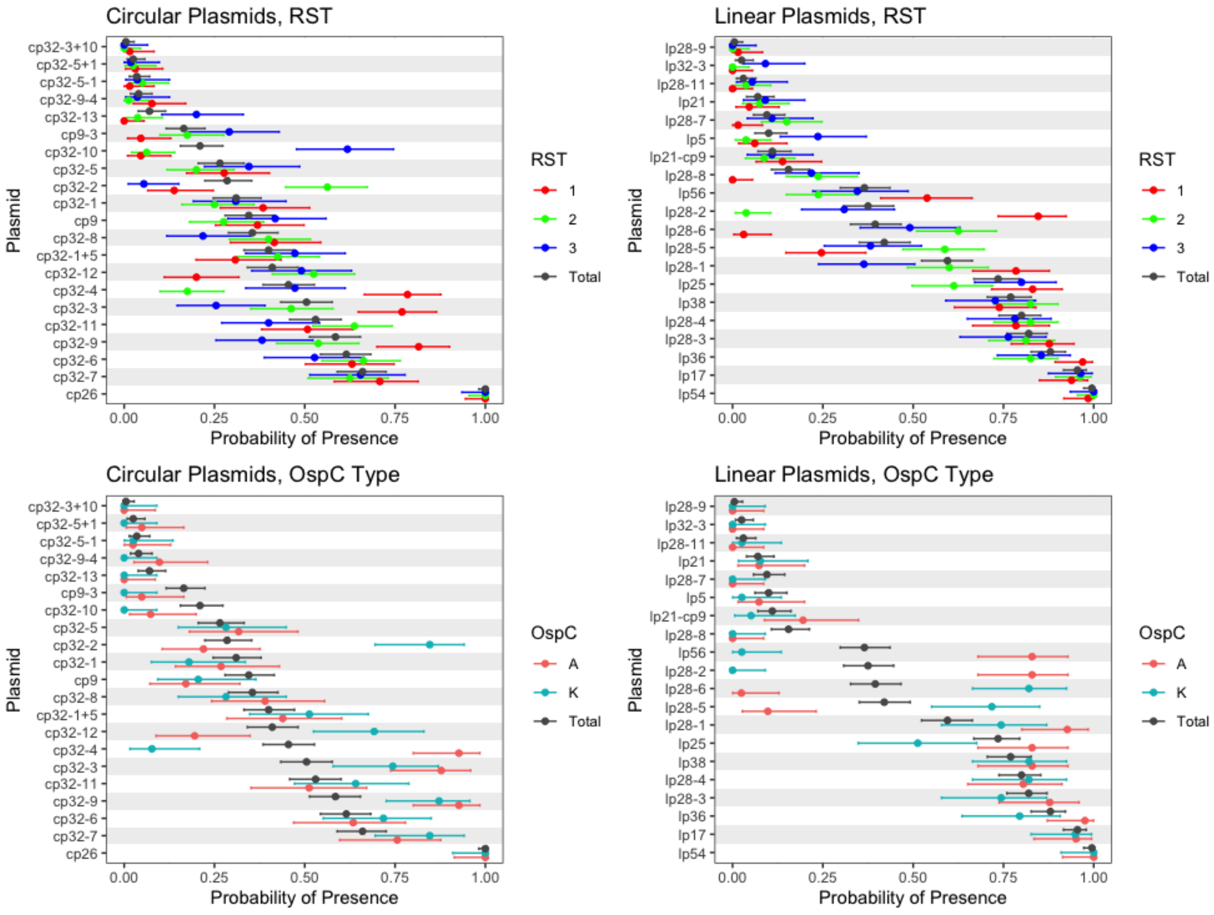


**Figure S2: Genotype-conditional probabilities of plasmid presence.** Point estimates are shown with 95% Clopper-Pearson confidence intervals. Analysis highlights RST-specific variations, as well as differences between OspC types A and K, compared to the aggregate probabilities. Most notably, cp32-2, cp32-4, lp28-2, and lp28-5 exhibit the largest genotype-specific differences in presence patterns.

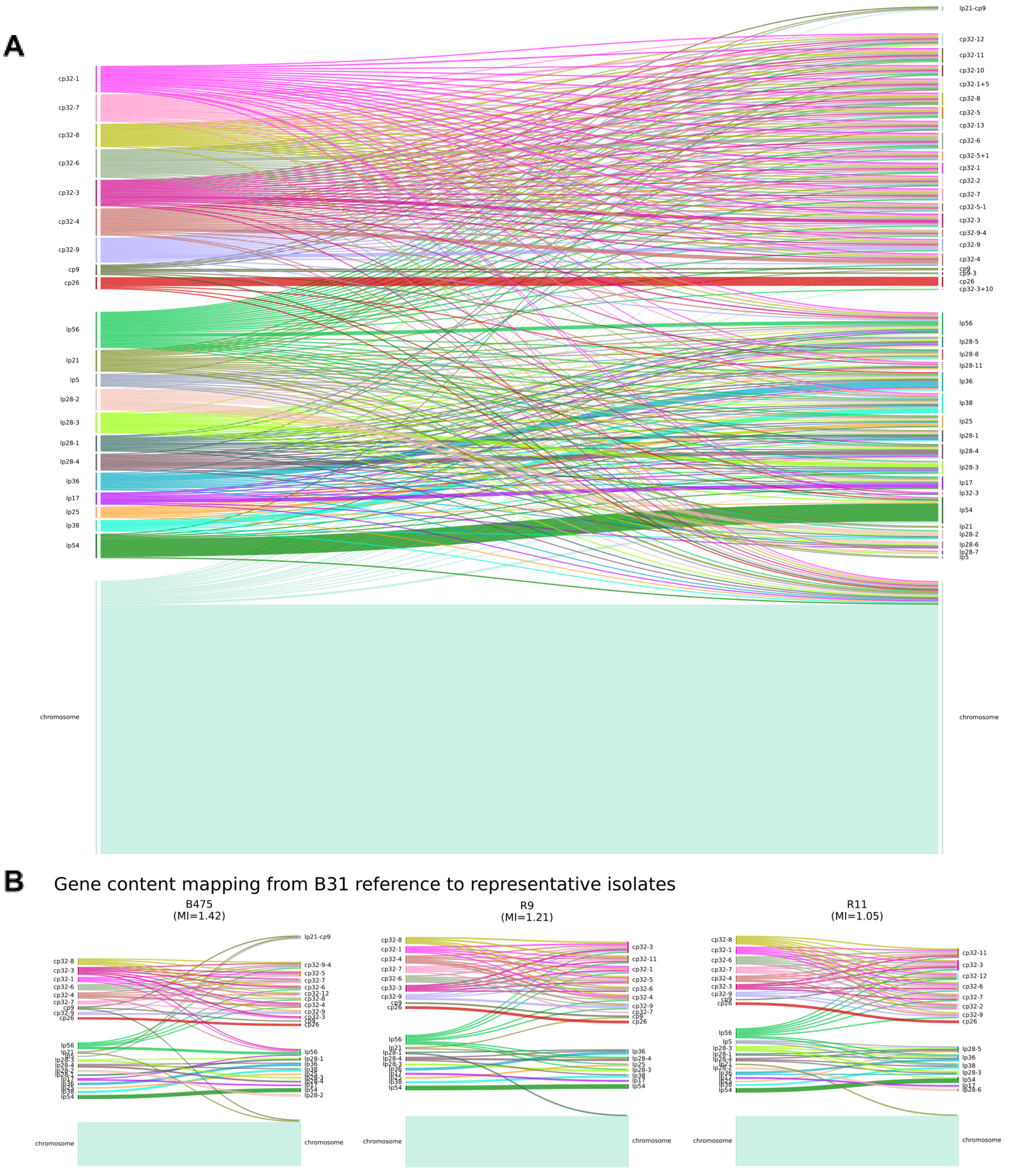
A
cp32-1
cp32-7
cp32-8
cp32-6
cp32-3
cp32-4
cp32-9
cp9
cp26
lp56
lp21
lp5
lp28-2
lp28-3
lp28-1
lp28-4
lp36
lp17
lp25
lp38
lp54
chromosome
lp21-cp9
cp32-12
cp32-11
cp32-10
cp32-1+5
cp32-8
cp32-5
cp32-13
cp32-6
cp32-5+1
cp32-1
cp32-2
cp32-7
cp32-5-1
cp32-3
cp32-9-4
cp32-9
cp32-4
cp9
cp9-3
cp26
cp32-3+10
lp56
lp28-5
lp28-8
lp28-11
lp36
lp38
lp25
lp28-1
lp28-4
lp28-3
lp17
lp32-3
lp54
lp21
lp28-2
lp28-6
lp28-7
lp5
chromosome
B
Gene content mapping from B31 reference to representative isolates
B475
(MI=1.42)
R9
(MI=1.21)
R11
(MI=1.05)
chromosome

C

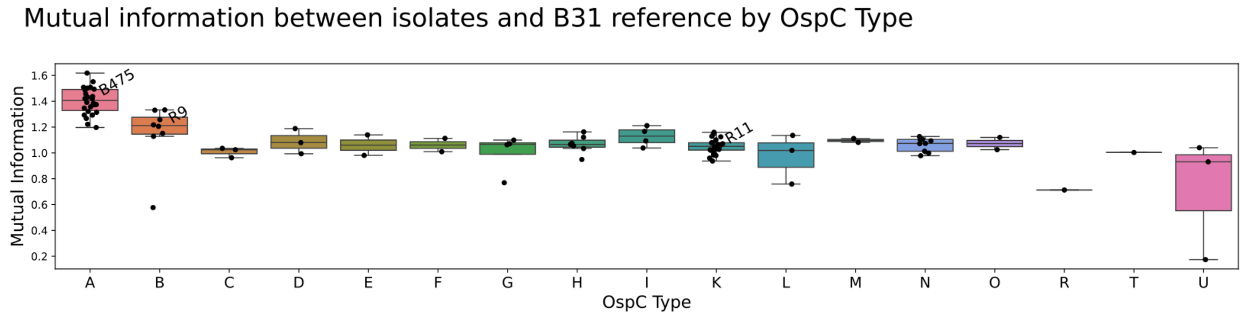


**Figure S3: Regions of homology between B31 reference isolate and assemblies.** (A) Sankey plot illustrates where genes are found in LRAs (right column) compared to the B31 reference genome (left column). Replicons in both columns are first grouped by replicon type (chromosome, linear plasmid, circular plasmid, fusion plasmid [bottom to top]), and then ordered by entropy (uncertainty) such that the most entropic (chaotic) replicons are at the top and the least entropic replicons are at the bottom. For the purpose of visualization, connecting curve widths have been rescaled to be more similar in size; scaled size is proportional to number of genes according to scaled_width = (number_of_genes + avg_number_of_genes) / 2. (B) Sankey plots showing homology between the B31 reference and representative individual strains for isolates corresponding to median normalized mutual information in OspC Types A, B, and K. (C) Normalized Mutual Information between isolated genomes compared to reference genome, broken out by OspC Type. The isolates from panel B are labeled.

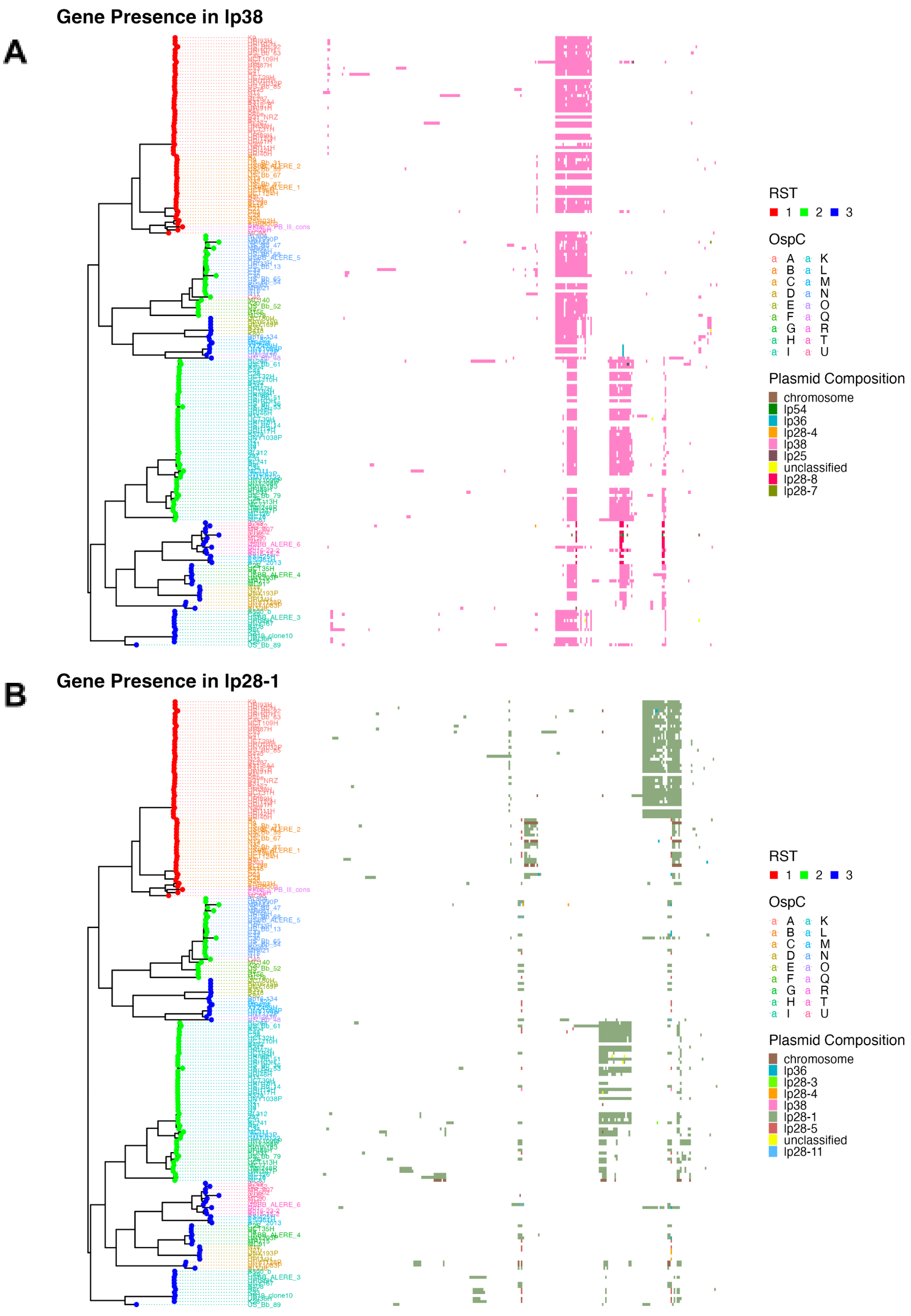


**Figure S4: Phylogenetic structure and gene presence at the plasmid level for (A) lp38 and (B) lp28-1.** Phylogenetic tree assembled from core genome sequences is shown at left. Tips are colored by RST and tip labels are colored by OspC type. To the right of the phylogenetic tree, a matrix shows orthologs present in each isolate that are encoded on that replicon. Columns are ordered according to hierarchical clustering based on Euclidean distances.

**A**

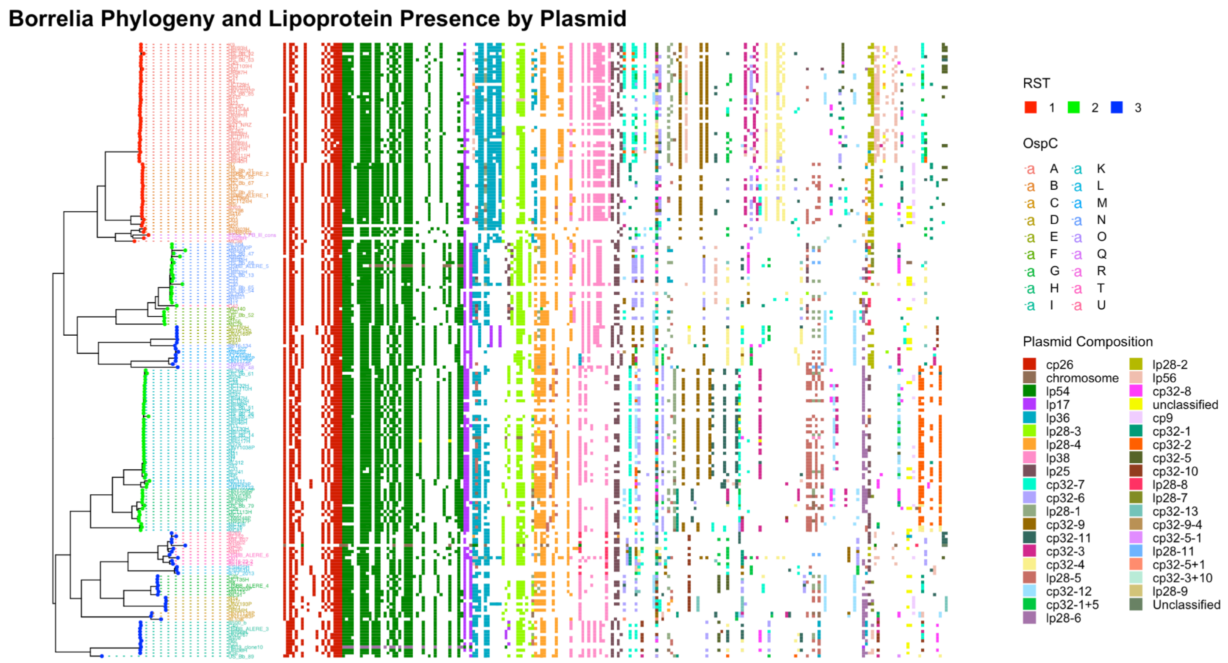


**Figure S5: Phylogenetic structure and plasmid occupancy patterns of Bb lipoprotein orthologs.** (A) Phylogenetic tree assembled from core genome sequences is shown at left. Tips are colored by RST and tip labels are colored by OspC type. To the right of the phylogenetic tree, lipoprotein orthologs present in each isolate are shown in a matrix. Columns are ordered based on best-hit plasmid frequency (i.e. if a gene is most commonly found on lp54, it goes with the lp54 set, and because lp54 is a core plasmid, it is the leftmost plasmid). The plasmid encoding the lipoprotein ortholog is annotated by color. As with Figure 2, we only include orthologs that are present in at least 10% of isolates. (B) Enlarged version of Figure 2 (attached as Supplemental File 1), with individual gene groups labeled. Note: chromosomal orthologs were removed. Note also that ESI26H lacks a fragment of lp54 but most individual genes are absent, suggesting that the plasmid is in the process of being lost.

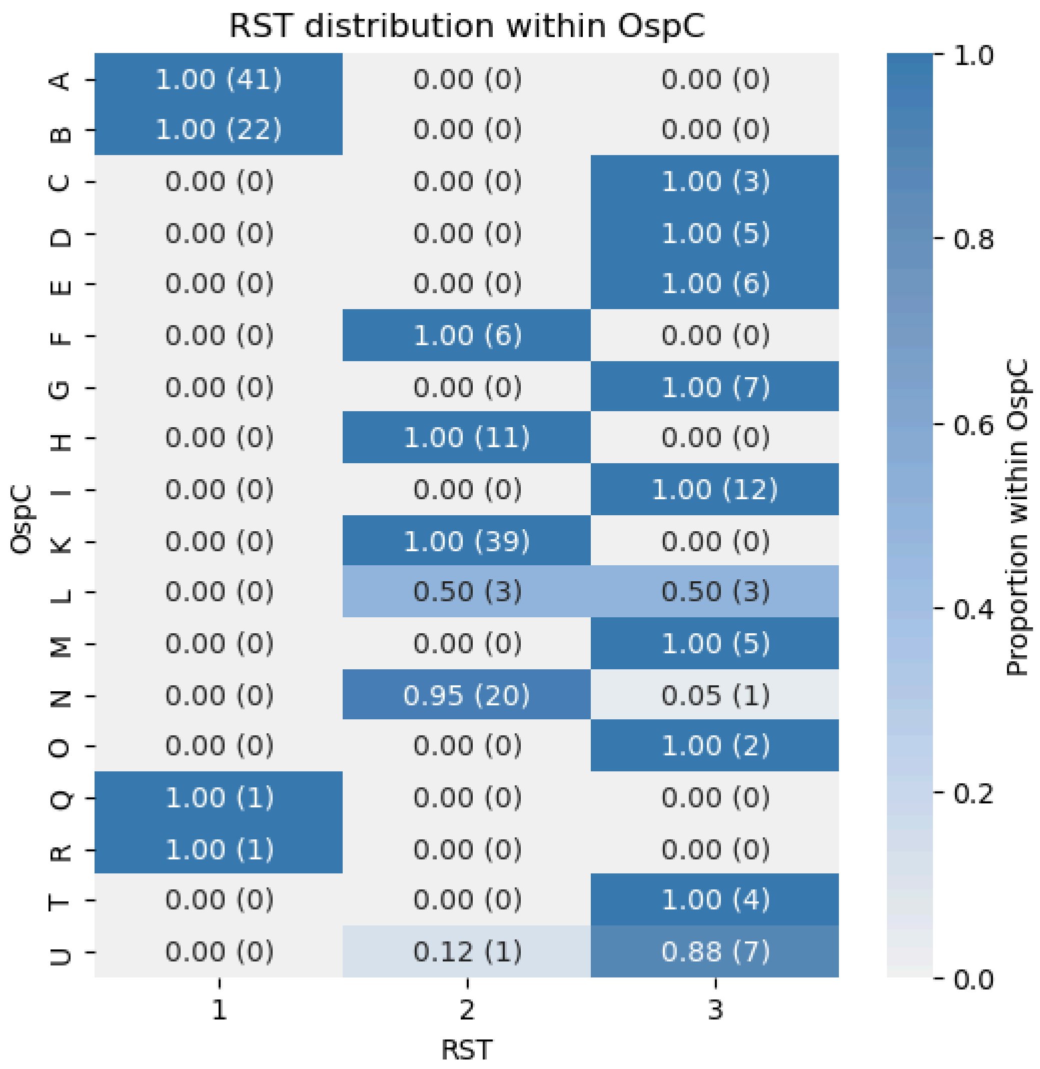


**Figure S6: Distribution of RST types within each OspC category.** Cells of the heatmap are annotated with both proportion and isolate count. Almost all OspC types are contained within a single RST, with the exception of OspC L which partitions evenly between RST2 and RST3, and OspC N and U which each have a single diverging strain.

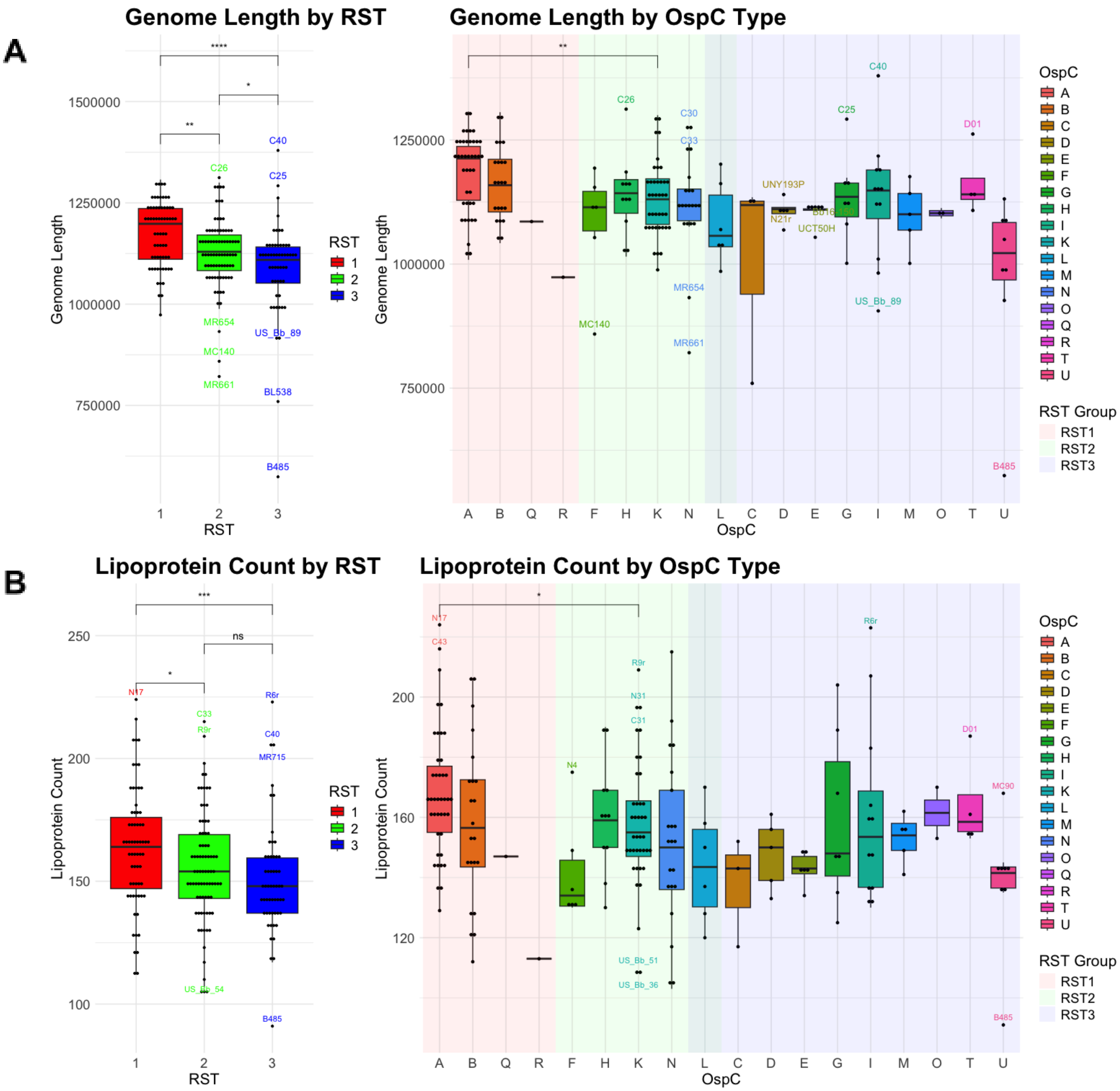
A
Genome Length by RST
Genome Length
RST
1
2
3
****
**
*
C40
C25
C26
MR654
MC140
MR661
US_Bb_89
BL538
B485
Genome Length by OspC Type
OspC
RST Group
RST1
RST2
RST3
A
B
Q
R
F
H
K
N
L
C
D
E
G
I
M
O
T
U
C30
C33
MR654
MR661
UNY193P
N21r
UCT50H
D01
B
Lipoprotein Count by RST
Lipoprotein Count
***
ns
N17
C33
R9r
R6r
C40
MR715
US_Bb_54
B485
Lipoprotein Count by OspC Type
N17
C43
N4
N31
C31
US_Bb_51
US_Bb_36
MC90

C

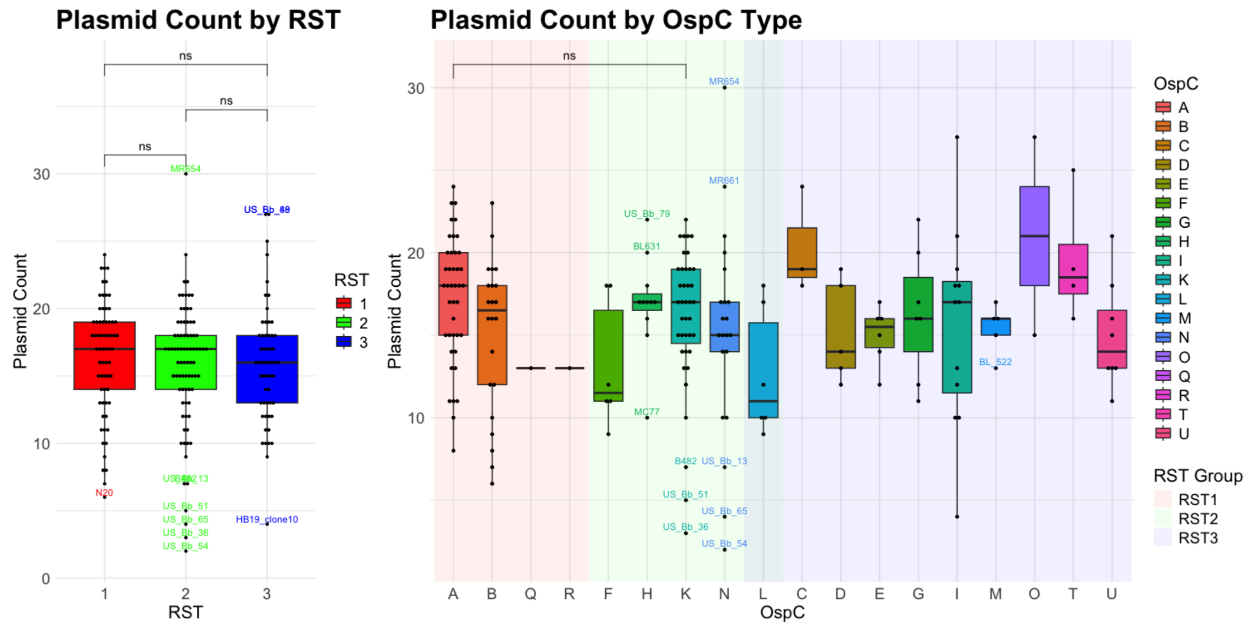


**Figure S7: (A) Genome size, (B) lipoprotein count, and (C) plasmid count distributions across RST and OspC types.** Pairwise significance testing (Wilcoxon rank-sum test) is shown between all RST types, and between OspC types A and K. Not significant - ns; *: P < 0.05; ** - p < 0.01; *** p < 0.001 ; **** p < 0.0001. OspC types are sorted by their best-fit RST category. Most notably, OspC types A and K differ significantly with respect to genome length and lipoprotein count, but not plasmid count. This also holds true for RST1 and RST2, and RST1 and RST3.

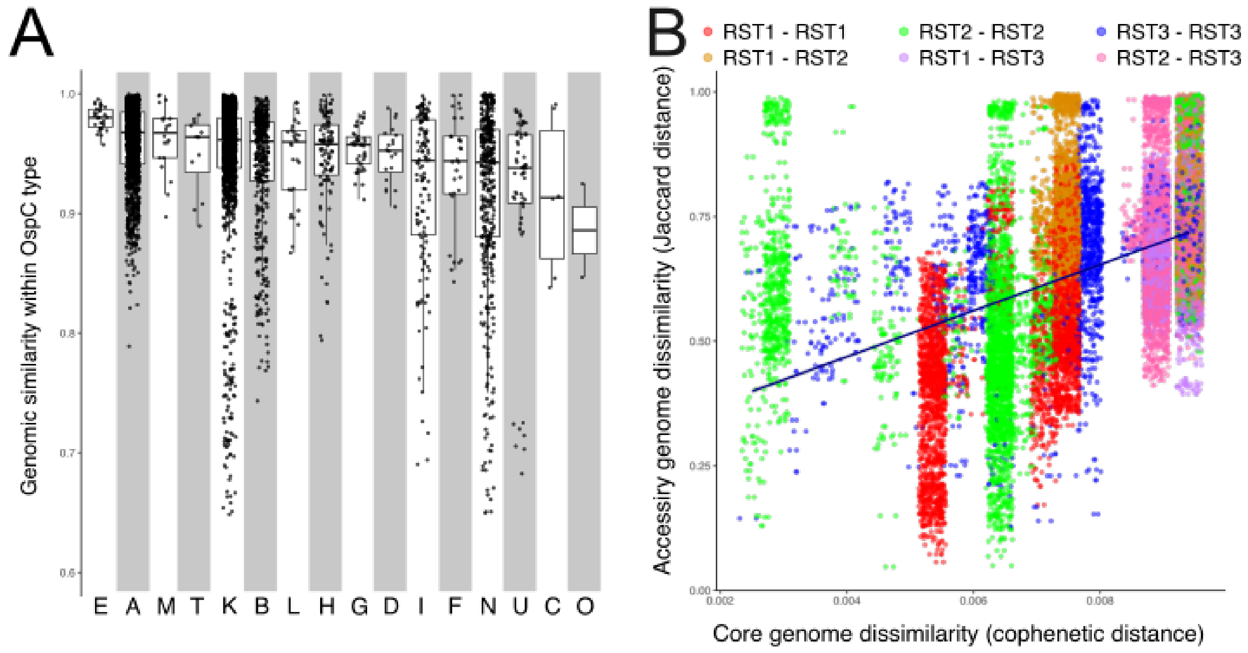


**Figure S8: Genome divergence and similarity in Bb population.** (A) Boxplots depicting genomic similarity via full average nucleotide identity (ANI) within each OspC type. (B) Scatterplot showing the projection of core (cophenetic distances) versus accessory (Jaccard distances) genome divergence, with a fitted linear regression.

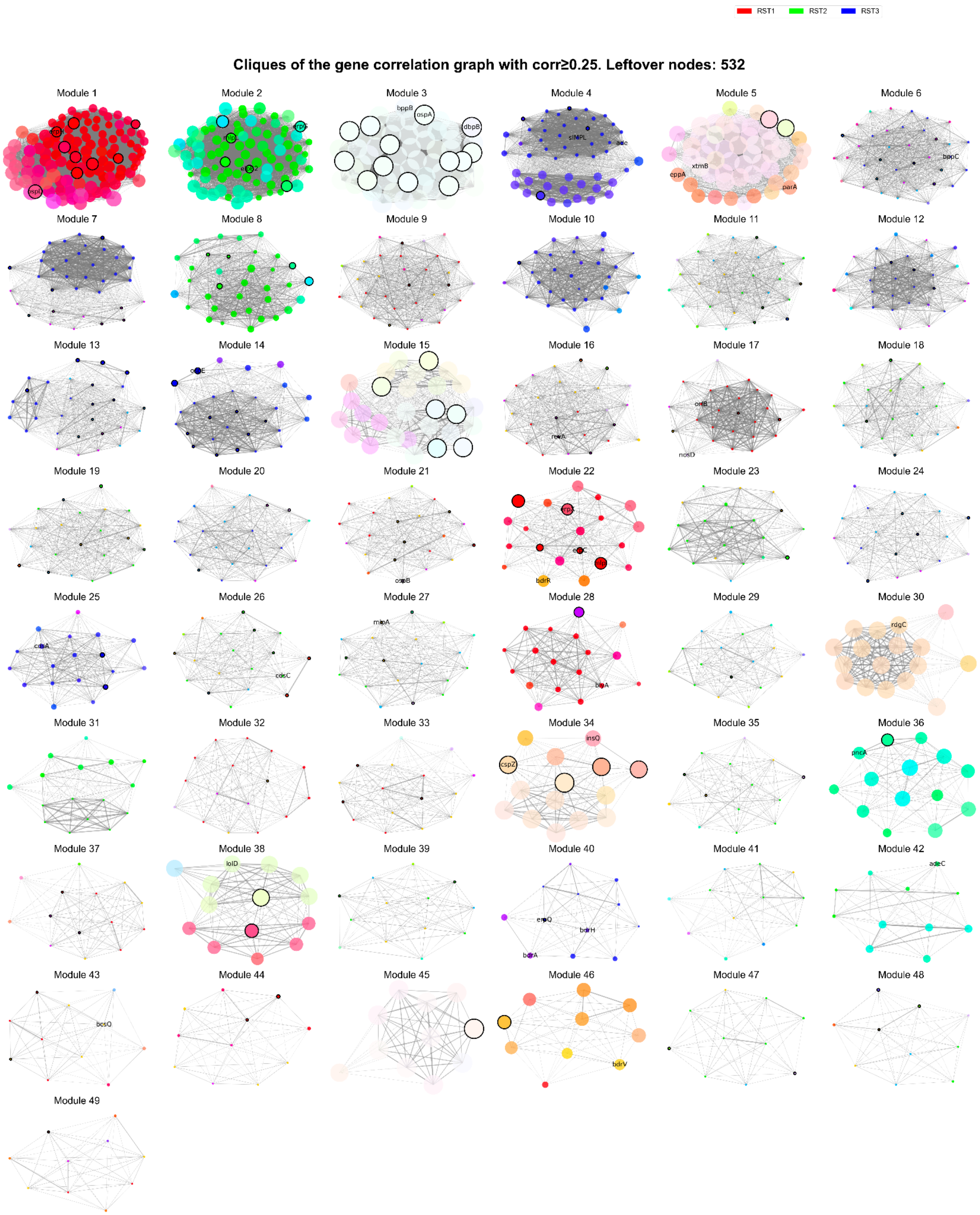


**Figure S9: Modular analysis reveals 49 distinct modules within the *Bb* genome.** Cliques were found iteratively upon a subgraph of the full correlation network where genes present in only one isolate were dropped and the minimum correlation threshold was set to 0.25. For clarity, only cliques with 10 or more nodes are displayed. Node colors are determined by the RST to RGB colormapping as in Figure 4. Surface lipoproteins are circled in black, and certain gene groups with sufficiently short names are annotated.

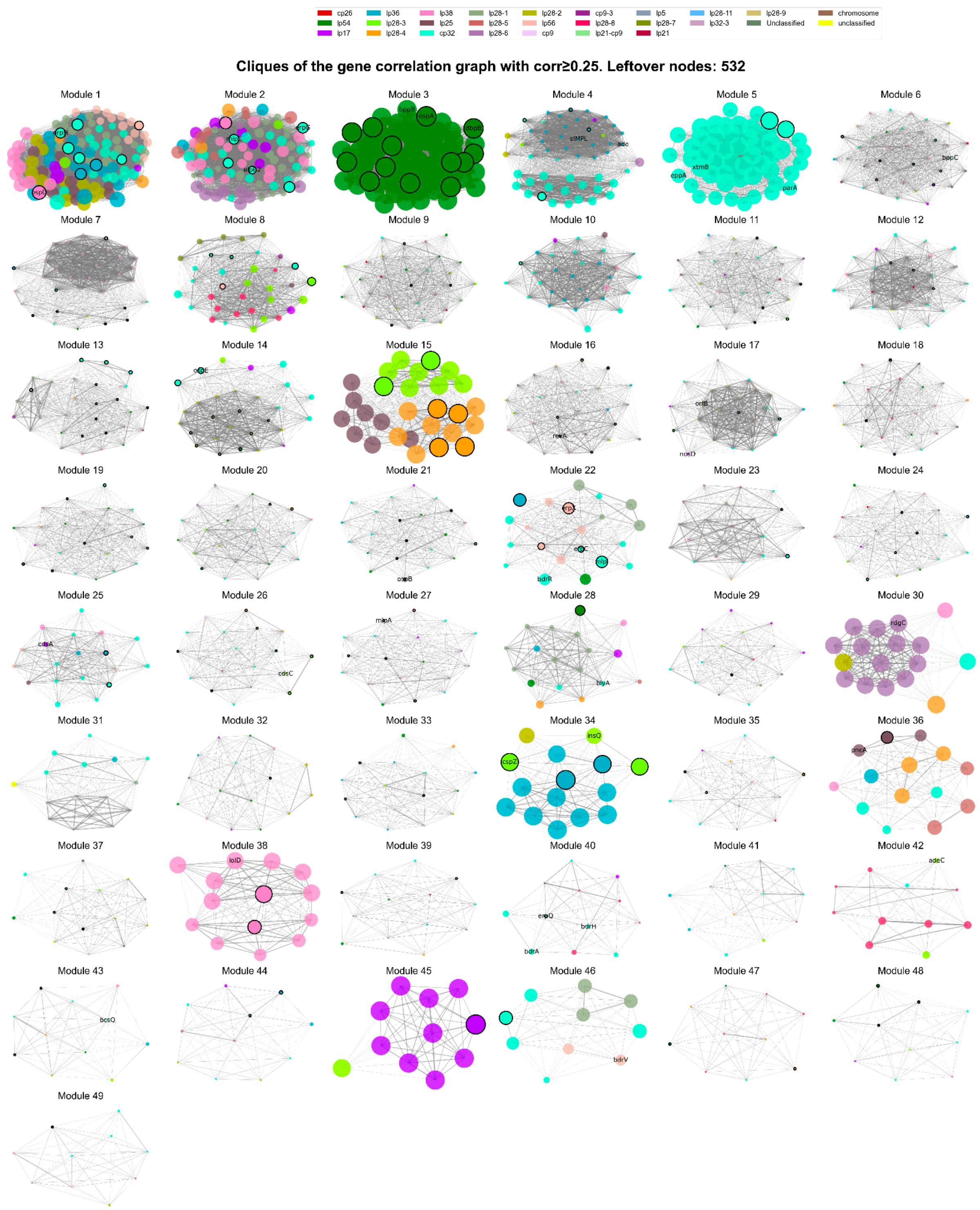


**Figure S10: Modular analysis colored by plasmid demonstrates structural connectivity of gene groups across replicons.** These modules are defined exactly as in Figure S8, with gene groups colored by the replicon they are most commonly found on. For visual clarity, gene groups belonging to the cp32 family share the same color. Surface lipoproteins are circled in black, and certain gene groups with sufficiently short names are annotated.

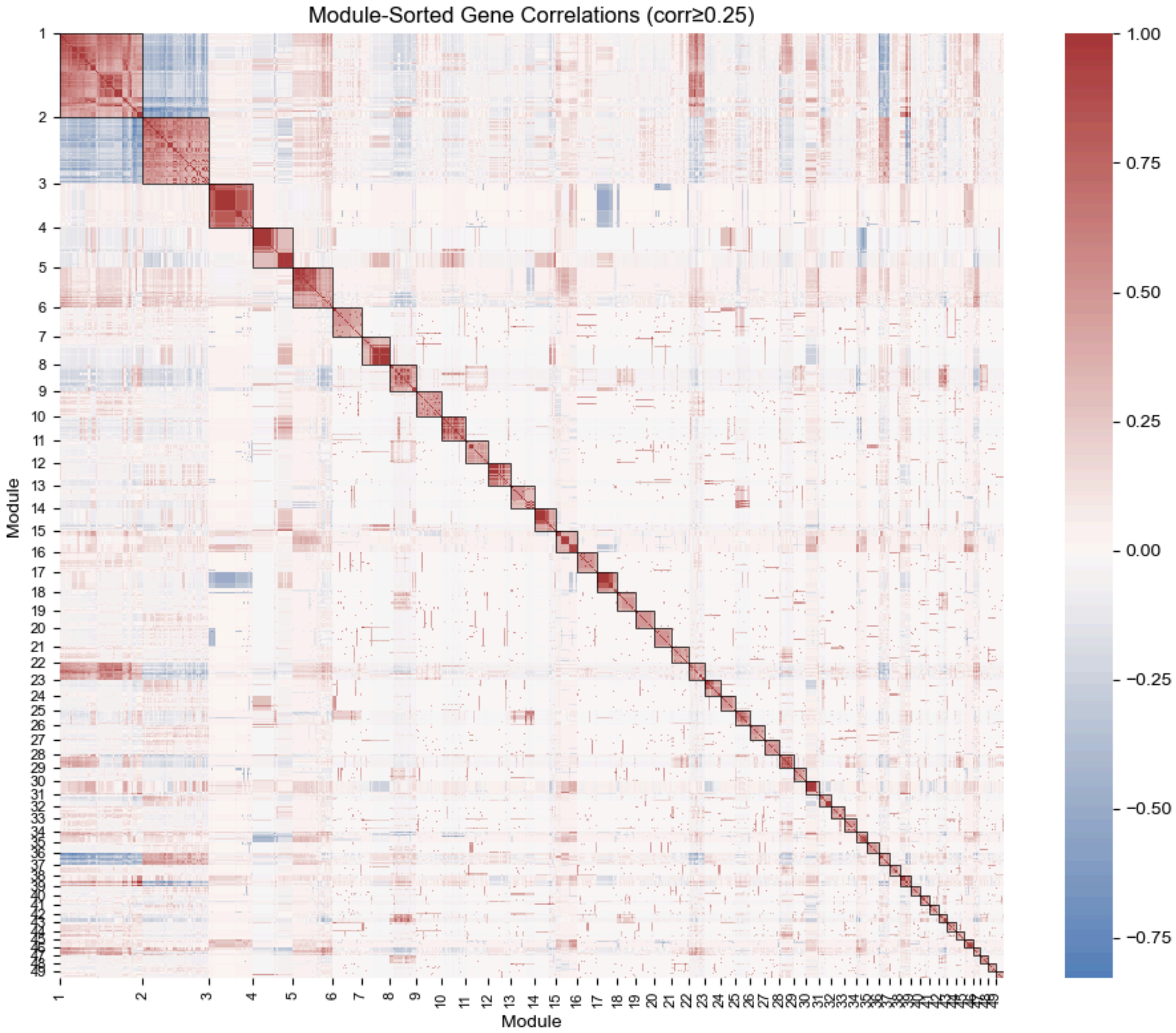


**Figure S11: Heatmap of gene correlations according to presence / absence patterns.** Represents Pearson's correlation between all pairs of genes. Genes are grouped together and labelled by module as in Figure S8. The strongest positive pairwise correlations occur along the diagonal, indicating that the iterative clique-finding methodology has partitioned the full correlation network into meaningful modules.

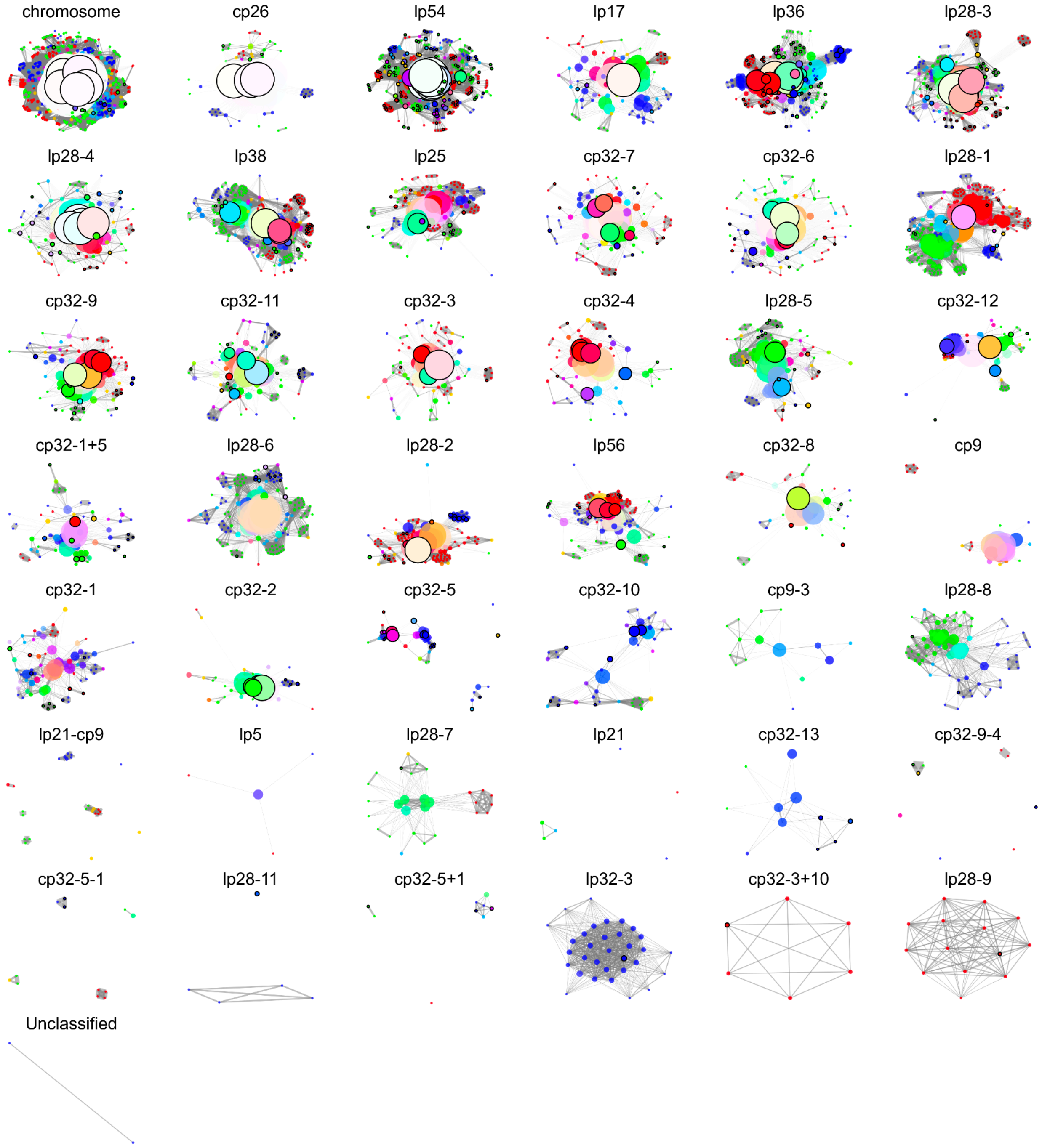


**Figure S12: Plasmid networks demonstrate differing gene compositions depending on RST type.** Genes were partitioned into independent subsets based on the plasmid they were most frequently found on across our isolates. Induced subgraphs were then constructed using positive correlations between nodes within each set. These modules are subgraphs of the pangenome correlation graph shown in Figure 4b. Surface lipoproteins are circled in black. Based on their gene content, various plasmids are more heavily weighted to certain RST types, which is indicative of evolutionary patterns in the *Bb* genome.

**Appendix of Supplemental Tables and Files**

**Table S1:** List of strains and isolate metadata

**Table S2:** Contingency table of isolate sources

**Table S3:** Summary of ortholog counts in the pangenome

**Table S4:** Hypergeometric test for lipoprotein presence in k-partite modules

**Table S5:** List of k-partite modules and gene content

**Table S6:** Contingency tables of genotypes with dissemination

**Supplemental File 1:** High-resolution version of Figure 2.

| | Human | Tick | Total |
|---|---|---|---|
| **Sequenced through LRA** | 182 | 1 | 183 |
| **Downloaded from GenBank** | 6 | 11 | 17 |
| **Total** | 188 | 12 | 200 |

**Table S2:** Contingency table of isolate sources.

| # Genes (% of isolates) | Merged paralogs pangenome | Unmerged paralogs pangenome | Sig. level | P Value | Test |
|---|---|---|---|---|---|
| Core (99%+) | 796 | 702 | *** | 4.60e-12 | proportions z-test |
| Soft core (95-99%) | 102 | 160 | ns | 5.53e-02 | proportions z-test |
| Shell (15%-95 %) | 506 | 897 | *** | 4.50e-12 | proportions z-test |
| Cloud (<15%) | 4614 | 5668 | ns | 6.31e-01 | proportions z-test |
| **Total** | **6018** | **7427** | *** | **3.47e-19** | $\chi^2$ **test** |

**Table S3:** Counts of ortholog groups in the pangenome, with and without splitting of paralogous gene families. The p-value reports the test of hypothesis that counts of split and unsplit paralogs are the same, using a z-test of proportions.

| RST | Disseminated | Localized | No Data | Total |
|---|---|---|---|---|
| **1** | 25 | 24 | 16 | 65 |
| **2** | 37 | 30 | 13 | 80 |
| **3** | 16 | 22 | 17 | 55 |
| **Total** | 78 | 76 | 46 | 200 |

| OspC | Disseminated | Localized | No Data | Total |
|---|---|---|---|---|
| **A** | 20 | 11 | 10 | 41 |
| **B** | 4 | 12 | 6 | 22 |
| **C** | 3 | 0 | 0 | 3 |
| **D** | 2 | 3 | 0 | 5 |
| **E** | 3 | 1 | 2 | 6 |
| **F** | 1 | 5 | 0 | 6 |
| **G** | 1 | 5 | 1 | 7 |
| **H** | 7 | 2 | 2 | 11 |
| **I** | 4 | 6 | 2 | 12 |
| **K** | 17 | 17 | 5 | 39 |
| **L** | 1 | 1 | 4 | 6 |
| **M** | 2 | 1 | 2 | 5 |
| **N** | 11 | 5 | 5 | 21 |
| **O** | 0 | 2 | 0 | 2 |
| **Q** | 1 | 0 | 0 | 1 |
| **R** | 0 | 1 | 0 | 1 |
| **T** | 0 | 1 | 3 | 4 |
| **U** | 1 | 3 | 4 | 8 |
| **Total** | 78 | 76 | 46 | 200 |

**Table S6:** Contingency tables of dissemination status with (A) RST and (B) OspC Type.
