## Supplementary figures and images for "Linkage blocks involving multiple replicons organize *Borrelia burgdorferi* strain variation and influence virulence in Lyme disease"

### SupplementaryFileS1

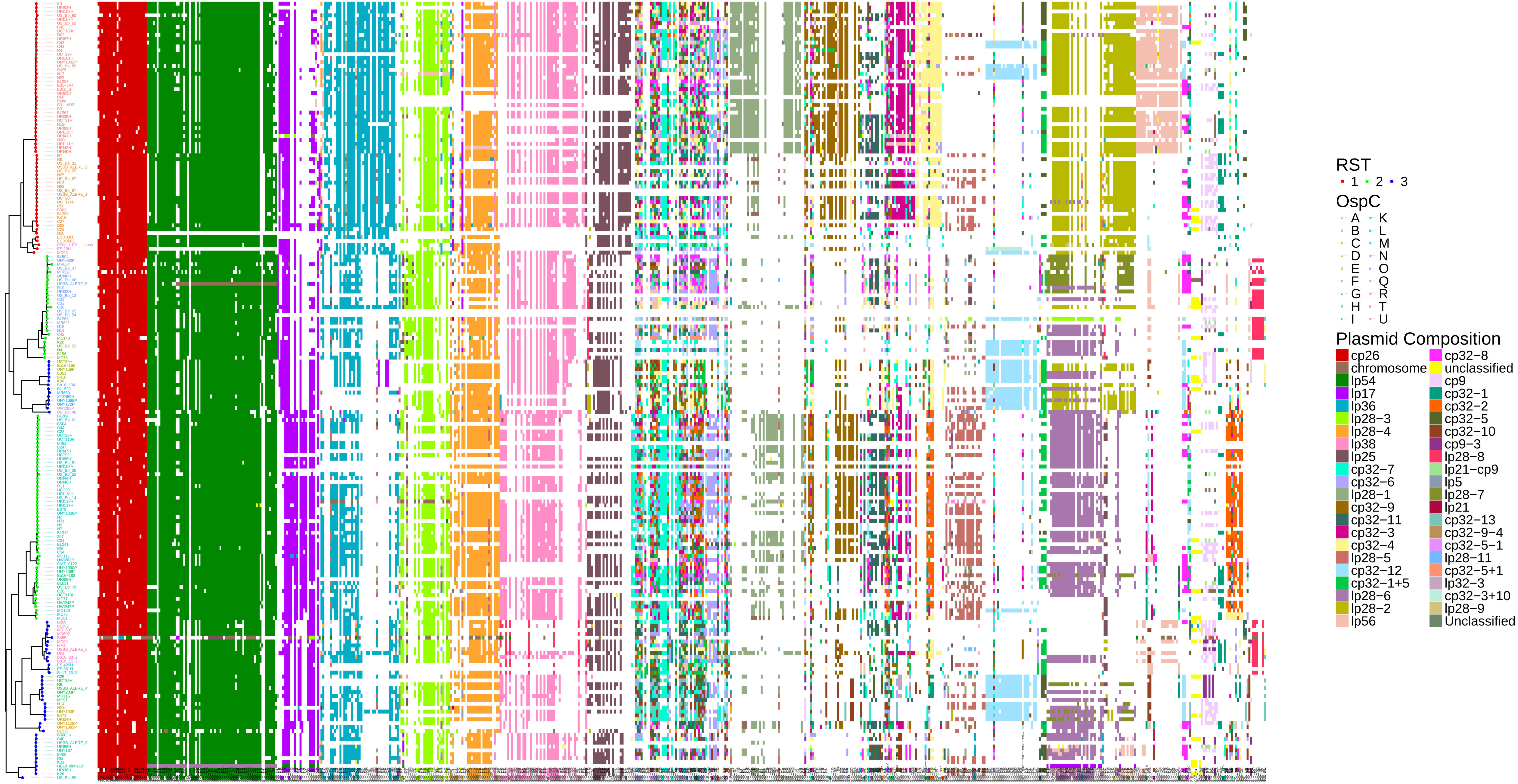
